# Low level of antioxidant capacity biomarkers but not target overexpression predicts vulnerability to ROS-inducing drugs

**DOI:** 10.1101/2023.01.17.524372

**Authors:** Jana Samarin, Piotr Fabrowski, Roman Kurilov, Hana Nuskova, Johanna Hummel-Eisenbeiss, Hannelore Pink, Nan Li, Vivienn Weru, Hamed Alborzinia, Umut Yildiz, Laura Grob, Minerva Taubert, Marie Czech, Michael Morgen, Christina Brandstädter, Katja Becker, Lianghao Mao, Ashok Kumar Jayavelu, Angela Goncalves, Ulrike Uhrig, Jeanette Seiler, Yanhong Lyu, Sven Diederichs, Ursula Klingmüller, Martina Muckenthaler, Annette Kopp-Schneider, Aurelio Teleman, Aubry K Miller, Nikolas Gunkel

## Abstract

Despite a strong rationale for why cancer cells are susceptible to redox-targeting drugs, such drugs often face tumor resistance or dose-limiting toxicity in preclinical and clinical studies. An important reason is the lack of specific biomarkers to better select susceptible cancer entities and stratify patients. Using a large panel of lung cancer cell lines, we identified a set of “antioxidant-capacity” biomarkers (ACB), which were tightly repressed, partly by STAT3 and STAT5A/B in sensitive cells, rendering them susceptible to multiple redox-targeting and ferroptosis-inducing drugs. Contrary to expectation, constitutively low ACB expression was not associated with an increased steady state level of reactive oxygen species (ROS) but a high level of nitric oxide, which is required to sustain high replication rates. Using ACBs, we identified cancer entities with a high percentage of patients with favorable ACB expression pattern, making it likely that more responders to ROS-inducing drugs could be stratified for clinical trials.

## Introduction

It has long been postulated that modulating the levels of reactive oxygen species (ROS) is a promising means to kill cancer cells without compromising normal cells [1]. This concept hinges on the idea that tumors have a more oxidized redox homeostasis, which is adjusted to execute cancer hallmarks like survival, metastasis, vascularization and proliferation[2]. This redox status causes collateral damage to biomolecules, and tumors must compensate by upregulating antioxidant proteins. Cancer cells are therefore thought to be highly dependent on these antioxidant proteins and their finely tuned expression and activity levels. As such, inducing a shift in a tumor’s redox homeostasis by reducing antioxidant capacity or increasing ROS production should alter the critical ROS balance of cancer cells, resulting in cell cycle arrest or cell death while sparing non-cancerous cells [3]. There is increasing interest in redox-modulating drugs, not only because ROS-induced cell death and cancer selectivity offer a conclusive rationale for anticancer therapy, but also because many clinical drugs and development candidates induce elevated ROS levels, even though their mode of action is not directly connected to redox regulating targets [4].

In spite of their promise, drugs directly targeting ROS homeostasis have found only limited success in clinical trials, compared to other classes of anticancer agents [5]. This raises the question whether cancer cells may not be generally more susceptible to ROS stress than cells from normal tissues, resulting in a smaller therapeutic window than anticipated. However, another clearly important reason for the unimpressive performance of ROS-inducing drugs is that the field currently lacks a biomarker concept that meets the complexity and functional redundancy of the tumor redox system and which enables the selection of patients who will benefit most from redox-targeted therapies [5]. With such a concept established, both preclinical and clinical studies can be designed to reduce unnecessary and expensive failures.

In order to facilitate the discovery of new redox-targeting molecules and to support the development of clinical candidates, we set out to identify molecular markers with the highest possible predictive power for response to ROS-inducing drugs. We hypothesized that the prediction of drug efficacy is more robust when combining multiple markers. We challenged a panel of 31 non-small cell lung cancer (NSCLC) cell lines with a novel, potent and selective inhibitor of thioredoxin reductase 1 (TXNRD1), recently developed by our group [6], for its central role in redox regulation. We identified a set of 15 genes with a strong correlation to drug sensitivity, which we named ACBs (for antioxidant capacity biomarkers). With the ACB biomarker set established in our NSCLC cell line panel, we were able to predict drug response in a collection of NSCLC patient derived xenograft (PDX) models. A cross-entity analysis revealed that the majority of cancers, including lung adenocarcinoma (LUAD), expressed a mostly unfavorable biomarker profile, consistent with the limited success of redox targeting drugs in clinical trials. We also identified cancer entities where a substantial percentage of patients should show response redox-targeting drugs.

In the course of this study, we discovered characteristics of drug-sensitive cells, which contradict the current thinking on ROS homeostasis in cancer cells. We found that both resistant and sensitive cells, although they express distinctly different levels of ACBs, have established a similar, low steady-state redox status, challenging the notion that elevated basal ROS levels are a hallmark of sensitivity towards drug-induced ROS. Unexpectedly, we found that sensitive cells express high levels of nitric oxide (NO), which they depend on for cell proliferation and, to a certain extent, to compensate for the lack of enzymatic ROS buffer capacity. Sensitive cells remain robustly vulnerable to drug-induced ROS as they stably silence ACB genes, by mechanisms partly elucidated in this study.

## Materials and Methods

### Synthesis of DKFZ-682

(*S*)-(+)-prolinol (98 %) and sodium aurothiomalate(I) (99.9 % trace metal basis) were obtained from Alfa Aesar, carbon disulfide (analytical reagent grade) was obtained from Fisher Scientific UK, potassium hydroxide (≥85 %) was obtained from Carl Roth, Germany, deionized water was used from in-house supply. All chemicals were used as received without further purification. NMR spectra were recorded on Bruker 400 MHz instrument at 298.1 K. High resolution mass spectrometry was recorded on a Bruker ApexQe FT-ICR instrument, (Department of Organic Chemistry, University of Heidelberg). Elemental analyses were performed on a vario Micro Cube by the “Microanalysis Laboratory”, Institute of Organic Chemistry, University of Heidelberg.

### [Au^I^((*S*)-prolinoyl)dtc] (DKFZ-682)

A 250 mL two-necked round bottom flask equipped with an overhead stirred was charged with (*S*)-(+)-prolinol (2.94 g, 2.87 mL, 29.07 mmol, 1.2 equiv) dissolved in 40.0 mL of water (*aqua dest.*). To this solution was added KOH (1.90 g, 33.92 mmol, 1.4 equiv) and the mixture was stirred at 24 °C for 10 min. Then CS_2_ (2.58 g, 2.05 mL, 33.92 mmol, 1.4 equiv) was added dropwise with a syringe and the resultant mixture was stirred at 24 °C for 2 h assuming a complete formation of the corresponding dithiocarbamate ligand. In a separate flask sodium aurothiomalate (ATM) (9.45 g, 24.23 mmol, 1.0 equiv.) was dissolved in 40.0 mL of water (*aqua dest.*) using a ultrasonic bath and occasional gentle heating in a water bath (50 °C). After allowing to cool to 24 °C this yellow ATM solution was added in one portion to the before-mentioned ligand solution. An immediate appearance of a bright yellow precipitate indicates complex formation. The resultant suspension was stirred at 24 °C for 18 h. The suspension was then filtered through a sintered glass Büchner funnel (ø=6 cm, h=6 cm; Porosity 4) and the yellow solid was thoroughly washed with deionized water (800 mL), transferred into a 250 mL round bottom flask, resuspended in a minimum amount of deionized water and lyophilized over 72 h. The pure product was obtained as a pale yellow solid in a yield of 8.76 g (11.74 mmol, 97 %) (*CAVE:* The compound proved to be highly active/cytotoxic and the material obtained from lyophilization easily form aerosols/dusts. It is suggested to handle this pure (undissolved) material with appropriate safety measures such as wearing FFP3 masks when working outside a ventilated fumehood!).

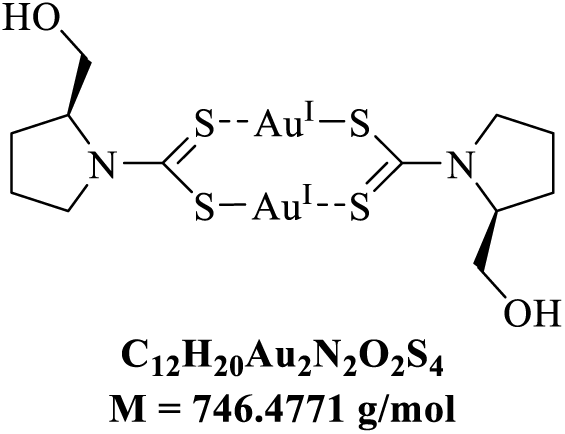

### Inhibitory capacity of DKFZ-682

Recombinant human glutathione reductase (GSR), human thioredoxin reductase isolated from placenta (TXNRD1), and recombinant, selenocysteine-lacking human TXNRD1(U498C) were produced as described previously [7]. The inactivation of GSR was measured in the GSSG reduction assay [8]. The GSR assay system contained 20.5 mM KH2PO4, 26.5 mM K_2_HPO_4_, 1 mM EDTA, 200 mM KCl, and pH 6.9 as buffer as well as 100 µM NADPH, 4 nM GSR, and 1 mM GSSG. Inactivation of TXNRD1 and TXNRD1(U498C) was measured in the DTNB and TXN reduction assay as follows: DTNB substrate assay: TXNRD1 was diluted in 0.1 M Tris, 1 mM EDTA pH 7.4, 500 µM NADPH to a concentration of 90 nM and aliquoted on a drug dilution plate. Test compounds were added to the first aliquot and serial dilutions were prepared in enzyme mix. From each concentration 75 µL were pipetted in a Greiner F-bottom 96 well plate in triplicate. 15 min after the addition of the test compound, the reaction was started by the addition of 25 µL DTNB solution. The reaction was monitored by determination of absorption at 420 nm for 30 min in a FLUOstar OPTIMA ELISA reader in 20 sec intervals. The enzyme activity (OD 420 nm/min) was calculated from the initial, linear part of reaction. Fluorescent TXN assay: A commercial kit, including cloned mammalian TXNRD1, TXN1 and fluorescent insulin was used (IMCO, Stockholm, Sweden, Cat No. FkTRXR-03-STAR) according to the instructions of the supplier. Fluorescence (485/520 nm) was monitored in a FLUOstar OPTIMA ELISA reader in 20 sec intervals. Both assays were conducted in 47.4 mM KH_2_PO_4_, 52.6 mM K_2_HPO_4_, pH 7.4, 100 µM (TXN assay) or 200 µM (DTNB assay) NADPH. In the DTNB and TXN reduction assays the final concentration of redox enzymes and substrates were as follows: 16 nM (TXN assay) or 4 nM TXNRD1 (DTNB assay); 2.5 µM (TXN assay) or 3 µM TXNRD1 (U498C) (DTNB assay), and 50 µM TXNC72S (TXN assay) or 3 mM DTNB (DTNB assay), respectively.

Half-maximal inhibition of the respective enzymes was determined by incubating the enzymes (GSR, TXNRD1 and TXNRD1(U498C)) for 3 min with NADPH (100 or 200 μM) followed by the addition of different inhibitor concentrations at 25 °C using a Tecan Infinite M200 plate reader (Tecan). All assays contained a negative control (no compound = 0 % inhibition), and a positive control (no substrate = 100 % inhibition). Compounds were dissolved in DMSO (Auranofin) or 200 mM cyclodextrine (DKFZ-682). After monitoring the baseline, the reaction was started by adding the respective substrate and ΔA/min was monitored at 340 or 412 nm, respectively.

### Cell culture

The human NSCLC cell lines purchased from ATCC were cultured in RPMI-1640, high-or low glucose DMEM or DMEM/F12 media (Gibco, Sigma-Aldrich), containing 10 % fetal bovine serum (FBS-12A, Capricorn scientific), 100 U/mL penicillin, and 100 µg/mL streptomycin (Sigma-Aldrich) at 37 °C in a 5 % CO_2_ atmosphere. For some cell lines (H1793, H23, H1693, H1568) RPMI-1640 medium contains additional 10 mM HEPES, 1 mM sodium pyruvate, and 2.5 g/L glucose. Cells were splitted by incubation with trypsin-EDTA for 5 min at 37 °C. CRISPRa cell lines were selected for puromycin (sgRNAs) and blasticidin (dCas9-VP64 construct) every 2-3 weeks for 72 h to make sure that only cells that carry both insertions survive. Therefore, 4 µg/mL puromycin and 25 µg/mL blasticidin were added to the according cell culture medium.

The human AML cell lines available from the German Collection of Microorganisms and Cell Cultures (DSMZ) were maintained in suspension cultures in the recommended culture media at 37 °C in a 5 % CO_2_ atmosphere. Specifically, HEL, NOMO1, U937, GDM1, KG1, EOL1, MOLM14, NB4, HL60, and THP1 were cultured in RPMI-1640 supplemented with 10 % FBS, MOLM16, ME1, PL21, HNT34, and KASUMI1 in RPMI-1640 supplemented with 20 % FBS, M07E in RPMI-1640 with 20 % FBS and 10 ng/mL GM-CSF and OCIM1 in Iscove’s MDM with 10 % FBS. All the media contained 100 U/mL penicillin and 100 µg/mL streptomycin. Prior to seeding for experiments, cells were harvested by centrifugation (3 min at 300 g at room temperature).

### Chemical compounds

Chemical compounds used in this study are listed in Supplementary Table S1.

### CellTiter-Blue assay and determination of EC50

Cells were seeded in 96-well plates (Greiner, F-bottom) at a density of 5,000 to 15,000 cells/well dependent on cell line and incubation time. 24 h later test compounds were added (all concentrations in triplicate) without medium change. After further incubation for indicated time period, viable cells were quantified in a FLUOstar OPTIMA ELISA reader (550/590 nm) about 2 h after CellTiter-Blue staining (Promega, Cat No. G8081). Mean values +/-standard deviations (SD) were calculated. EC50 were calculated from dose response curves by GraphPad Prism (log(inhibitor) versus response, variable slope (four parameters)).

### Monitoring of TXNRD1 activity in cells with TRFS-Green

TRFS-Green was synthetized according to Zhang et al. [9]. Cells were seeded in Greiner V-bottom 96-well plates (7,500 cells/well) in Fluorobright DMEM medium. Inhibitors were added 4 h later in triplicates at the indicated concentrations, instantly followed by the addition of TRSF-Green to a final concentration of 10 µM. After mixing with a multi-channel pipette, plates were centrifuged at 200 g for 5 min. Fluorescence (450/520 nm) was monitored in a FLUOstar OPTIMA ELISA reader in 2 min intervals over a period of 6 h. For drawing dose response curves, the increase of fluorescence during the linear phase (e. g. 100 min) was calculated for each well. Mean values +/-SD were plotted.

### Measurement of ROS levels in cytoplasm and mitochondria

The effect of drugs on roGFP2-Orp1 oxidation was quantified as described [10] in the NSCLC cell line H838 stably expressing roGFP2-Orp1 (with or without the mitochondrial targeting sequence). The day before the measurement cells were seeded into a black clear-bottomed 96-well imaging plate (Falcon, Cat No. 353219) at a density of 20,000 cells/well in 200 µL Fluorobrite medium. A non-transduced control was included on the same plate for background subtraction. In order to obtain the fluorescence intensity values for a fully oxidized and reduced probe, control wells were treated with 2 mM diamide or 10 mM DTT for 15 min at 37 °C. After the entire plate was measured for 8 cycles in a CLARIOstar fluorescence plate reader, BMG Labtech (which allows the simultaneous detection of the two excitation maxima of roGFP2 (400 nm and 485 nm) when emission is monitored at 520 nm), 22 µL of 10x concentrated drug was added and measurement continued for up to 340 min. The readout of the roGFP2 measurement was expressed as the degree of sensor oxidation (OxD, see equation in [11]). All treatments were performed in technical triplicate seeded in different quadrants of the imaging plate, to avoid position effects.

### Detection of ROS/RNS by flow cytometry

For detection of ROS/RNS level 150,000 cells/well/1 mL (exception: 80,000/well in case of H661 and H1299 cell lines) were seeded in 12-well plate. The next day, cells were treated with DMSO (unstained control), 5 µM CM-H2DCFDA (Invitrogen^TM^, Thermo Scientific, Cat No. C6827), 1:250 OxiVision^TM^ Green peroxide sensor (AAT Bioquest, Cat No. 11506, powder was solved in 200 µL DMSO) or 5 µM DAF-FM Diacetate (Invitrogen^TM^, Thermo Scientific, Cat No. D23844) at 37 °C for 30 min; 1:400 DAX-J2™ PON Green (AAT Bioquest, Cat No. 16317) at 37 °C for 60 min. Then cells were detached with trypsin, washed, spun down 5 min at 1200 rpm and finally each cell pellet was resuspended in 500 µL PBS with 1 % FBS and analysed by the flow cytometer Guava easyCyte 14HT (Luminex). The fluorescence of all above mentioned dyes was analyzed in the Green-B channel (excitation 488 nm, emission 512/18 nm) while counting 10,000 cells per sample. For ROS Brite^TM^ 570 (10 µM, AAT Bioquest, Cat No. 16000) staining, cells were first detached with trypsin, washed and then incubated with ROS Brite^TM^ 570 at 37 °C. After 20 min cells were spun down 5 min at 1200 rpm and cell pellet was resuspended in 500 µL PBS with 1 % FBS and analysed by flow cytometry (Yellow-G channel: excitation 532 nm, emission 575/25).

Treatment: cells were co-incubated with OxiVision^TM^ Green peroxide sensor and drug; co-incubated with DAX-J2™ PON Green and drug; preincubated with drug, then detached with trypsin, washed and further incubated with ROS Brite^TM^ 570; preincubated with drug, then drug was removed and fresh medium with DAF-FM Diacetate was added. Stained but untreated cells were used as control.

### Growth rate

One day before the experiment, 1,000 cells (or 5,000 cells for H1944) were seeded on Corning® 96-well Flat Clear Bottom Black plates in five replicates for each day of measurements. The next day (day 0), cells were treated with 50 µM NO scavenger cPTIO (Sigma), 40 µM NO donor NOC-18 (Santa Cruz), or a combination of both. Control cells were treated with PBS. On day 0 and day 3, cell nuclei were stained with Hoechst 33342 and propidium iodide. The nuclei were counted with NYONE automatic microscope (Synentec). The proportion of nuclei positive for propidium iodide was similar for all treatments. Growth rates were calculated according to Hafner et al. [12] by using the formula:

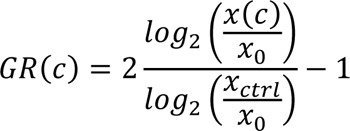

where x(c) is the treated nuclei count, x_ctrl_ is the control nuclei count and x_0_ is the nuclei count at the time of treatment (day 0).

### Detection of oxidized PRDX1 and PRDX3

For protein isolation, medium was removed from cells seeded in 6 cm dishes and cold thiol-block buffer (100 mM N-ethylmaleimide (NEM) in PBS) was added and incubated for 5 min on ice. Cells were lysed with 250 µL cold lysis buffer (1 % Triton X-100, 20 mM NEM in TBS (50 mM Tris, 150 mM NaCl, pH 7.4), complete protease inhibitor cocktail tablets (Serva, Cat No. 39101)) for 5 min on ice, sonicated and then centrifuged for 15 min at 12,000 x g at 4 °C. From each sample 100 µg protein was mixed with 4x Laemmli buffer (277.8 mM Tris-HCl pH 6.8, 26.3 % (w/v) glycerol, 2.1 % SDS, 0.01 % bromphenol blue (Na-salts), 40 mM NEM) for non-reducing condition and with 4x Laemmli buffer plus 20 % (v/v) 1M DTT for reducing condition. Samples were denatured for 5 min at 95 °C and separated on a denaturing gel (6 % stacking, 15 % separating). Proteins were transferred to a 0.2 µm nitrocellulose membrane using wet blot. Antibodies and respective dilutions are listed in Supplementary Table S2.

### Western immunoblotting

Cells were harvest using RIPA buffer (150 mM NaCl, 1 % (v/v) Nonidet P-40, 1 % (w/v) sodium deoxycholate, 0.1 % SDS, 50 mM Tris-Cl pH 8.0) supplemented with a protease inhibitor cocktail (Serva, Cat No. 39101.03), 100 U/mL benzonase (Merck) and PhosSTOP phosphatase inhibitor cocktail (Roche, Cat No. 04906837001). Protein concentrations were measured by Nanodrop (Thermo Fisher). Lysates were denatured by incubation with Laemmli sample buffer for 10 min at 95 °C and the proteins were separated by SDS-PAGE and transferred onto a nitrocellulose membrane using wet blot and incubate with gene specific antibodies (Supplementary Table S2). Immunoreactive proteins were visualized by the enhanced chemiluminescence detection system (Odyssey; LI-COR). The bands were quantified using the Licor image analyzer software. To correct for equal loading and blotting, all of the blots were redetected with antibodies directed against tubulin or GAPDH.

### siRNA knockdown of *NRF2* and *STAT*s

Small interfering RNA (siRNA) for the gene-specific inhibition of *NRF2*and *STAT*s were ordered from siTOOLS (Planegg, Germany). Transient transfection was performed using Lipofectamine® RNAiMAX Transfection Reagent according to the manufacturer’s instructions (Thermo Fisher Scientific, Germany).

### Multi-omics data

We obtained expression, proteomics, and metabolomics data from DepMap project portal (depmap.org). As an expression dataset we used log_2_ transformed (with pseudo-count of 1) RNA-seq TPM gene expression data for just protein coding genes (DepMap Public 20Q2) [13]. Proteomics dataset contained normalized protein expression data originating from mass spectrometry profiling [14]. Metabolomics dataset contained metabolite levels quantified using liquid chromatography–mass spectrometry [15]. Promoter methylation dataset (reduced representation bisulfite sequencing, file CCLE_RRBS_tss_CpG_clusters_20181022.txt.gz) included methylation values for clustered groups of CpGs within (−3,000, 2,000) nucleotides of the TSS for each gene [13].

Expression data of NSCLC PDX models were obtained from Champions Oncology. Units were converted to log_2_(TPM+1). TCGA-tumor expression data was downloaded from https://portal.gdc.cancer.gov/tcga. Units were converted to log_2_(TPM+1).

### Association analysis

We performed association analysis by calculating Pearson correlation between each individual feature (i. e. a vector of expression or metabolite levels for all tested cell lines) in each dataset and the vector of EC50 values. Results for each multi-omics set were plotted separately as a volcano plot using ggvolcano.corr function from BiocompR R package.

We then combined expression and proteomics correlation results by first calculating ranks for each gene separately in both sets using *p*-values from correlation tests (lower *p*-value → lower rank), and second calculating average rank. After that, we selected top genes that have the lowest average rank as the most important associations. We plotted heatmaps with expression levels for these top genes using ComplexHeatmap R package [16]. NRF2-dependency annotation was based on literature data [17], annotation for other transcription factors was obtained via Dorothea R package [18]. Taking the top 50 genes with the strongest associations to EC50, we used Dorothea to identify their potential TF regulators and selected the set of TFs that regulated at least two of the 50 genes.

### Compound similarity analysis

To assess the similarity between the biomarker profile of our compound and biomarker profiles of other compounds we looked at the associations between expression of the ACB set of 15 genes and AUC values in NSCLC adenocarcinoma cell lines that were screened in the CTRP [19] project.

For expression data, we used the same expression dataset from DepMap described above. AUC data for CTRP compounds was obtained using SummarizeSensitivityProfiles function PharmacoGx R package (with the option “published”) [20].

First, we calculated correlations between expression of each gene (from top 75 genes list) and drug response values. So, for each tested compound we got a 15-element vector with correlation coefficients for each of 15 ACB genes. Then we calculated the Euclidean distance between such vector for our compound and the vectors for all other compounds. Therefore, we got a measure of similarity, between our compound and all other compounds with similar biomarker profiles got smaller distances, while compounds with different profiles got larger distances. We plotted the heat map with corresponding correlation vectors for 50 compounds with smallest distances.

### Lable free proteome preparation

NSCLC cells were lysed in 1 % SDC buffer (1 % SDC, 100 mM Tris pH 8.5, 40 mM CAA and 10 mM TCEP), incubated on ice for 20 min, boiled at 95 °C, sonicated for 10 min on a Biorupter plus and heated again for 5 min at 95 °C as describe previously [21]. Proteins in the sample were digested with LysC (1:100) for 2 hours followed by Trypsin (1:100) for overnight at 37 °C. To the peptides 5x volume Isopropanol/1 % TFA was added and vortexed to stop the digestion. The peptides were de-salted on equilibrated styrenedivinylbenzene-reversed phase sulfonated (SDB-RPS) StageTips, washed once in isopropanol/1 % TFA and twice with 0.2 % TFA. Purified peptides were eluted with 60 µl of elution buffer (80 % v/v ACN, 1.25 % w/v NH4OH). The dried elutes were resuspended in MS loading buffer (2 % ACN, 0.1 % TFA) and stored at −20 °C until MS measurement.

### Data dependent acquisition (DDA)-PASEF measurement

Liquid chromatography was performed with a nanoElute (Bruker Daltonics Inc, Bremen, Germany) coupled online to a hybrid TIMS quadrupole TOF mass spectrometer (Bruker timsTOF Pro) via a CaptiveSpray nano-electrospray ion source and samples were measured in ddaPASEF mode. Samples (200 ng of peptide) were loaded onto a 25-cm reversed-phase column with 75 µM diameter, 1.7 particle size, and 120 A pore size (Aurora column, IonOpticks, Australia). Peptides were separated in 100 min active gradient at a flow rate of 250 nl min−1. Mobile phases buffer A and buffer B were 0.1 % formic acid (FA) and 99.9 % ddH2O and 0.1 % FA, 97.9 % ACN, and 2 % ddH2O respectively. Buffer B was linearly increased from 2 % to 12 % in 60 min, followed by an increase to 20 % in 30 min and a further increase to 30 % in 10 min, before increasing to 85 % for 10 min and holding that for additional 10 min. For the calibration of ion mobility dimension, TIMS elution voltages were calibrated linearly to obtain the reduced ion mobility coefficients (1/K0) using three Agilent ESI-Low Tuning Mix ions (m/z 622, 922 and 1222). For sample injection, the dda-PASEF windows scheme was recorded from m/z 150 to 1700 and 10 PASEF MS/MS scans per topN acquisition cycle. The precursor ion intensity threshold was 1000 a.u. and threshold for PASEF MS/MS ions was 10.000 a.u. TIMS functioning at Scan range 100-1700 m/z, Ramp Time 100 ms, Duty cycle 100 %, Cycle time 100.00 ms and Spectra Rate 9.43 Hz.

### LC MS/MS data processing

MS raw files were processed using Maxquant [22] version 1.6.1.17 supported by Andromeda search engine. The data was searched for proteins and peptides using a target-decoy approach with a reverse database against Uniprot Human (version 2021) Fasta file with a false discovery rate of less than 1 % at the levels of protein and peptide. No changes were made to the enabled default settings such as oxidized methionine (M), acetylation (protein N-term), and carbamidomethyl (C) as fixed modification and Trypsin as enzyme specificity. A maximum of 2 missed cleavages were allowed, and a minimum peptide length of seven amino acids set. The proteins were assigned to the same protein groups if two proteins could not be discriminated by unique peptides. The label-free quantification was performed using the MaxLFQ algorithm [23] and match between run feature was enabled for identification of peptide across runs based on mass accuracy and normalized retention times. For label free protein quantification minimum ratio count was set to 2. The Maxquant output table was analyzed in Perseus [24], prior to the analysis contaminants marked as reverse hits, contaminants and only identified by site-modification were filtered out.

### Statistical analysis

Data are presented as mean ± SD or SEM. Statistical analyses were performed using GraphPad Prism 9 software.

## Results

### A selective TXNRD1 inhibitor as a tool compound to identify biomarkers associated with high and low sensitivity to ROS induction

Due to its central role in ROS scavenging and signaling pathways, TXNRD1 is a prototypic target for ROS-inducing cancer therapy [25]. The gold-complex auranofin is the most frequently used inhibitor of TXNRD1, however, its high, nonspecific reactivity with exposed cysteines in proteins and glutathione, its short half-life in biological liquids [26] and its limited selectivity for the TXN system [27] pose handicaps for its use as a chemical probe. Therefore, we decided to use the gold (I)-dithiocarbamate complex DKFZ-682 (compound 37 [6]), which is a similarly potent TXNRD1 inhibitor as auranofin, but is much less active on glutathione reductase (GSR) and less inhibited by serum albumin (Fig. 1A; Supplementary Fig. S1A).

**Figure 1.**
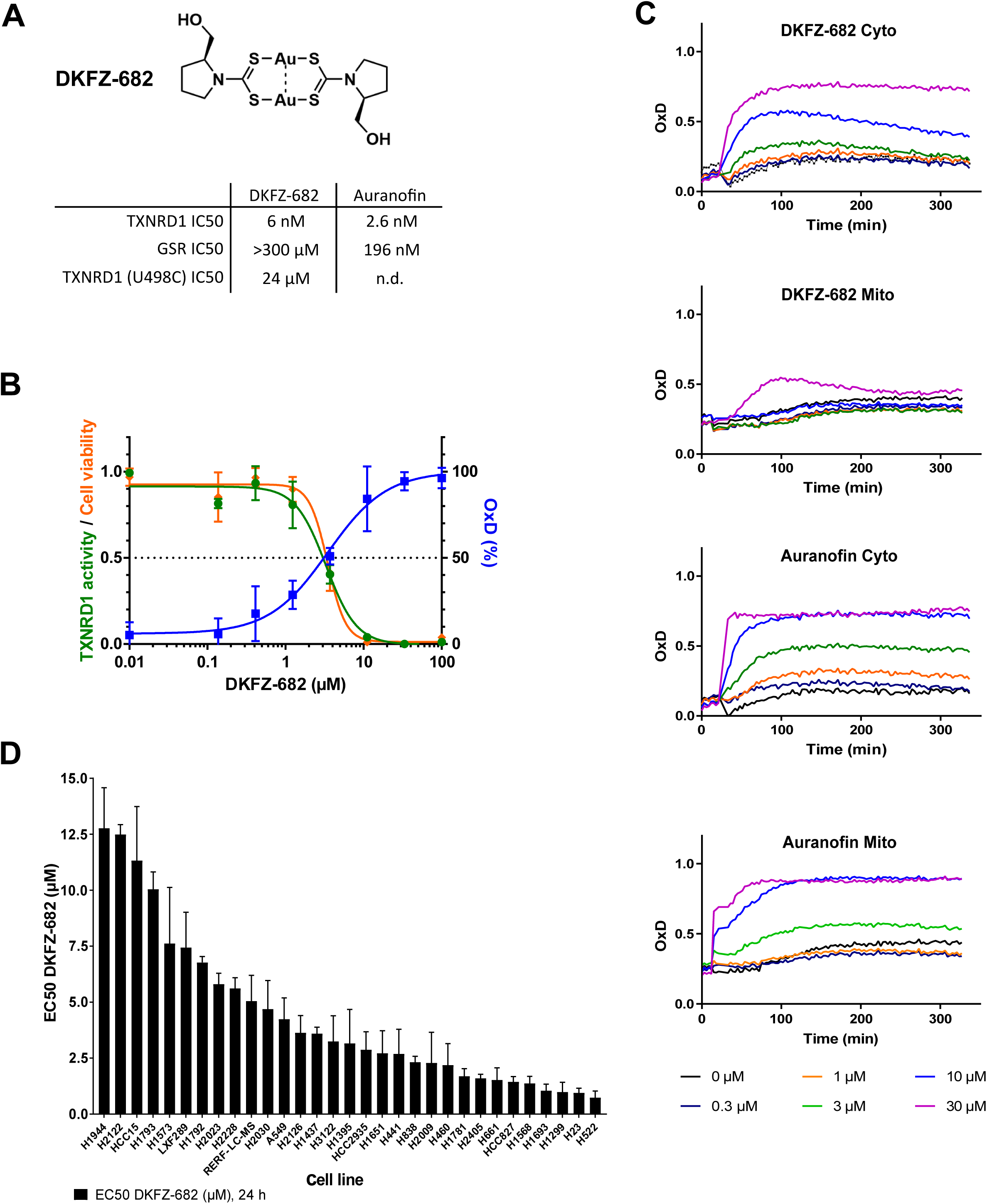
A selective TXNRD1 inhibitor as a tool compound to identify cells with high and low sensitivity to ROS induction. **(A)** Chemical structure of DKFZ-682, previously reported as gold (I)-dithiocarbamate (dtc) complex 37. Table shows 50 % inhibition values (IC50) of DKFZ-682 and auranofin on the main target, TXNRD1, and an off-target, GSR. Selectivity for the C-terminal selenocysteine in TXNRD1 is demonstrated by an increased IC50 value in the mutant protein TXNRD1 (U498C). **(B)** Dose-response curves representing DKFZ-682 activity characterized in H838 cells by measuring TXNRD1 enzymatic activity (TRFS-Green), roGFP probe oxidation, and cell viability. Orange line: cells were treated with a series of DKFZ-682 concentrations for 24 h. Cell viability was measured with CellTiter-Blue assay. Green line: cells were treated with TXNRD1 activity probe TRFS-Green and a series of DKFZ-682 concentrations. The signal intensity of the fluorescent reaction product was recorded over time. For each DKFZ-682 concentration, the initial enzyme velocity was calculated. Blue line: cells expressing ORP1-roGFP in the cytoplasm were treated with a series of DKFZ-682 concentrations. After 144 min, when the probe oxidation plateau was established, the oxidation (OxD, %) was calculated for each of the DKFZ-682 concentrations. All results are presented as dose-response curves. Data are presented as mean ± SD of three technical replicates. **(C)** H838 cells expressing either cytoplasmic (Cyto) or mitochondrial (Mito) roGFP2-Orp1 were treated with various concentrations of DKFZ-682 or auranofin. The fluorescence of oxidized and non-oxidized cytoplasmic or mitochondrial roGFP2-Orp1 were monitored for 300 min. OxD is the degree of probe oxidation with the data normalized to 1.0 (fully oxidized) defined by the signal from diamide (2 mM) and 0.0 (fully reduced) defined by the signal from DTT (10 mM). Data are presented as mean ± SD of three technical replicates. **(D)** Thirty-one NSCLC cell lines were treated with a concentration series of DKFZ-682 for 24 h and the cell viability was quantified by the CellTiter-Blue assay. EC50 values were determined from dose-response curves using GraphPad Prism. Bar diagrams represent mean ± SD of DKFZ-682 EC50 (µM) results from independent experiments performed in triplicates (n=2-6).

Cytotoxicity induced by DKFZ-682 directly correlates with cellular enzyme inhibition and the induction of disulfides, as demonstrated by the TXNRD-selective off-on fluorescent probe TRFS-green [9] and by the genetically encoded hydrogen peroxide (H_2_O_2_) probe roGFP2-Orp1 [28], respectively (Fig. 1B). Another important predictor of a future ROS-inducing drug’s safety and tumor selectivity is mitochondrial toxicity [29]. Interestingly, the mitochondrial ROS induced by DKFZ-682 can be suppressed if DKFZ-682 is formulated with sulfobutylether-β-cyclodextrin (Fig. 1C), an effect which cannot be achieved with auranofin [6]. Our results indicate that DKFZ-682 is a potent and selective TXNRD1 inhibitor, and therefore a highly suitable chemical probe to identify biomarkers for the response to ROS-inducing drugs.

With this tool compound in hand, we assembled a panel of 31 NSCLC cell lines with a wide range of sensitivities against DKFZ-682, both in 2D (Fig. 1D) and 3D culture conditions (Supplementary Fig. S1B), thereby providing sufficient resolution to identify predictive biomarkers and to investigate the molecular bases for the sensitivity against drug-induced ROS stress.

### Hallmarks of sensitivity to ROS scavenger targeted cancer therapy: Sensitive cells tend to have higher levels of steady state RNS

One concept for selective toxicity of ROS-inducing drugs in cancer cells assumes that sensitive cells have higher basal levels of ROS, which is further increased beyond a toxic threshold by drug treatment [1]. To validate this notion, we characterized basal and induced levels of various ROS species using an array of fluorescent dyes (Fig. 2A). In addition, we examined reactive nitrogen species (RNS) as those may contribute to ROS homeostasis [30] and, to the best of our knowledge, have not been examined in the context of drug sensitivity.

**Figure 2.**
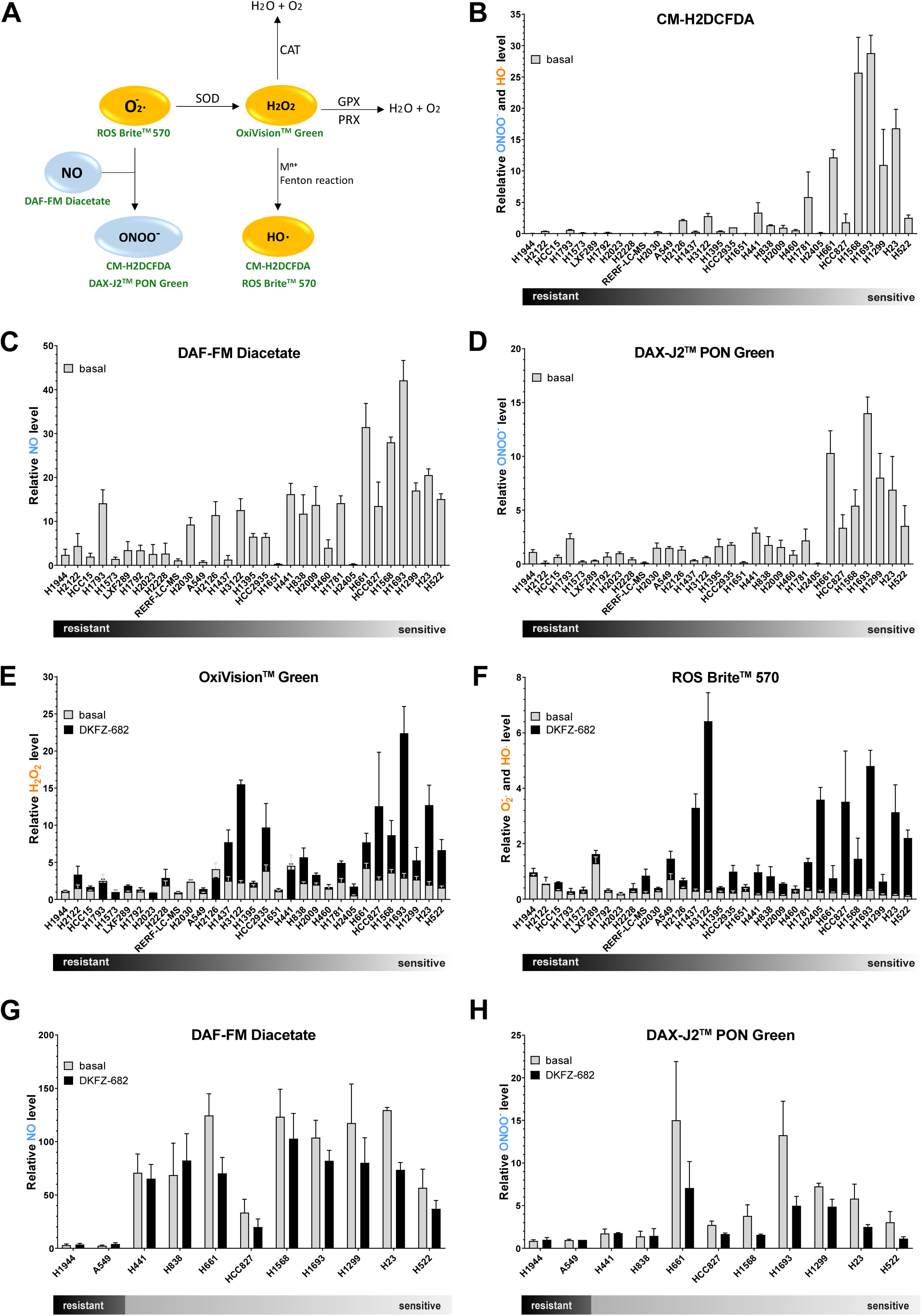
DKFZ-682 sensitive cells have higher levels of steady state RNS. **(A)** Simplified schematic representation of the main reactive oxygen and nitrogen species (ROS – orange; RNS - blue) and sensors (shown in green) for their detection. Superoxide dismutase (SOD) catalyzes the dismutation of superoxid anion (O_2_**^·^**^−^) to hydrogen peroxide (H_2_O_2_) and water (H_2_O). Superoxid anion may also react with nitric oxide (NO) to form peroxynitrite (ONOO^-^). Hydrogen peroxide may be degraded by catalases (CAT) and peroxidases (GPX, PRX) or it may be converted to the hydroxyl radical (HO^•^) via Fenton reaction (H_2_O_2_ oxidizes the reduced metal ion (M^n+^) to produce HO^•^). Sensors are nonfluorescent cell-permeant reagents and produce bright fluorescence upon ROS/RNS oxidation. The reagents have good selectivity to O_2_**^·^**^−^ (ROS Brite^TM^ 570), H_2_O_2_ (OxiVision^TM^ Green peroxide sensor), HO^•^ (CM-H2DCFDA, ROS Brite^TM^ 570), NO (DAF-FM Diacetate) and ONOO^-^ (DAX-J2™ PON Green, CM-H2DCFDA). **(B-H)** NSCLC cells were stained with CM-H2DCFDA **(B)**, DAF-FM Diacetate **(C, G)**, DAX-J2™ PON Green **(D, H)**, OxiVision^TM^ Green peroxide sensor **(E)** or ROS Brite^TM^ 570 **(F)** fluorescent dyes and analysed by flow cytometry. **(B)** Fluorescence in H2935 was set to 1 in each experiment. Bar diagrams summarize the data of independent experiments (n=3-8, error bars indicate SEM). **(C, D)** Fluorescence in H1944 was set to 1 in one experiment. The graphs summarize the relative data of independent experiments (n=3-6, error bars indicate SEM). **(E-H)** Cells were untreated or treated with DKFZ-682 (20 µM) for 30 min. Fluorescence in H1944 was set to 1 in one experiment. The graphs summarize the relative data of independent experiments (n=2-6, error bars indicate SEM).

Staining of unchallenged cells with CM-H2DCFDA, widely used to evaluate basal or induced cellular ROS levels, showed that sensitive cells exhibit higher fluorescent signals than resistant cells (Fig. 2B; Supplementary Fig. S2A). As CM-H2DCFDA is unable to discriminate between hydroxyl radical (HO^•^) and peroxynitrite (ONOO^-^) [31], we dissected the ROS/RNS profile of the cell panel with additional dyes.

DAF-FM diacetate and DAX-J2™ PON Green are fluorescent probes that predominantly detect NO [32] and ONOO^-^ [33], respectively. Staining of unchallenged cells showed higher levels of both RNS species in sensitive cells (Fig. 2C, 2D; Supplementary Fig. S2A, correlation with lnEC50: r=-0.62, *p*=0.0002 and r=-0.62, *p*=0.0002). Particularly for ONOO^-^, we noticed a distinct group of the 7 most sensitive cell lines which demonstrated nearly 10 times higher signals than a corresponding group of resistant cells. When we examined the ROS levels, using ROS Brite^TM^ 570 or OxiVision^TM^ Green peroxide sensor, we noted that the differences between resistant and sensitive cells were less pronounced. The high-ONOO^-^group of sensitive cells demonstrated 3-fold lower levels of superoxide (O_2_^•-^)/HO^•^ and 2-fold higher levels of H_2_O_2_ than their resistant counterparts (Fig. 2E, 2F; Supplementary Fig. S2A). The high correlation of CM-H2DCFDA and DAX-J2™ PON Green staining profiles (r=0.86, *p*<0.0001) suggests that our initial finding was based on ONOO^-^ level rather than on HO^•^.

It is interesting to note that the ROS/RNS profile of sensitive cells was inverted by drug treatment. ROS species like O_2_^•-^, HO^•^ or H_2_O_2_ were strongly increased (Fig. 2E, 2F) while NO and ONOO^-^ were reduced (Fig. 2G, 2H), most likely due to the reaction of H_2_O_2_ with NO [34]. Resistant cells, on the contrary, demonstrated a robust buffer capacity against drug-induced ROS or externally added H_2_O_2_ (Supplementary Fig. S2B).

The observations obtained with ROS-specific dyes were reflected in treatment-dependent dimer formation of cytoplasmic PRDX1 and mitochondrial PRDX3, both redox-sensitive peroxidases [35] (Supplementary Fig. S2C). Sensitive cells such as H661 and H838 do not have higher baseline oxidation of PRDX1/3 compared to resistant cells (e.g. H1944 or H1793), but have more oxidized PRDX1/3 in response to DKFZ-682 treatment. This shows that cells sensitive or resistant to ROS-inducing drugs have comparable basal ROS levels, and that sensitive cells are marked by a lower capacity to maintain redox homeostasis.

Taken together, these data demonstrate that sensitive cells have high levels of steady state RNS whereas basal ROS levels are inconspicuous. Our data also show that redox profiling with CM-H2DCFDA needs to be supplemented with more selective sensors in order to quantify steady state or induced ROS/RNS levels.

### High RNS status is favourable for cell proliferation

The data presented so far show that RNS rather than ROS levels are constitutively elevated in the sensitive cells, raising the question about the role of RNS in those cells. NO levels are likely to play a dominant role in this context as it is the rate-limiting precursor for the production of RNS [30]. NO reduction by the NO scavenger cPTIO caused strong inhibition of cell proliferation in sensitive, high-RNS cells (H1299, H23, HCC827), but not in resistant cells (A549, LXF289, H1944) (Fig. 3A). This effect could be rescued by the NO-donor NOC-18, suggesting that sensitive cells require high NO levels to support cell proliferation by a mechanism that has yet to be identified.

**Figure 3.**
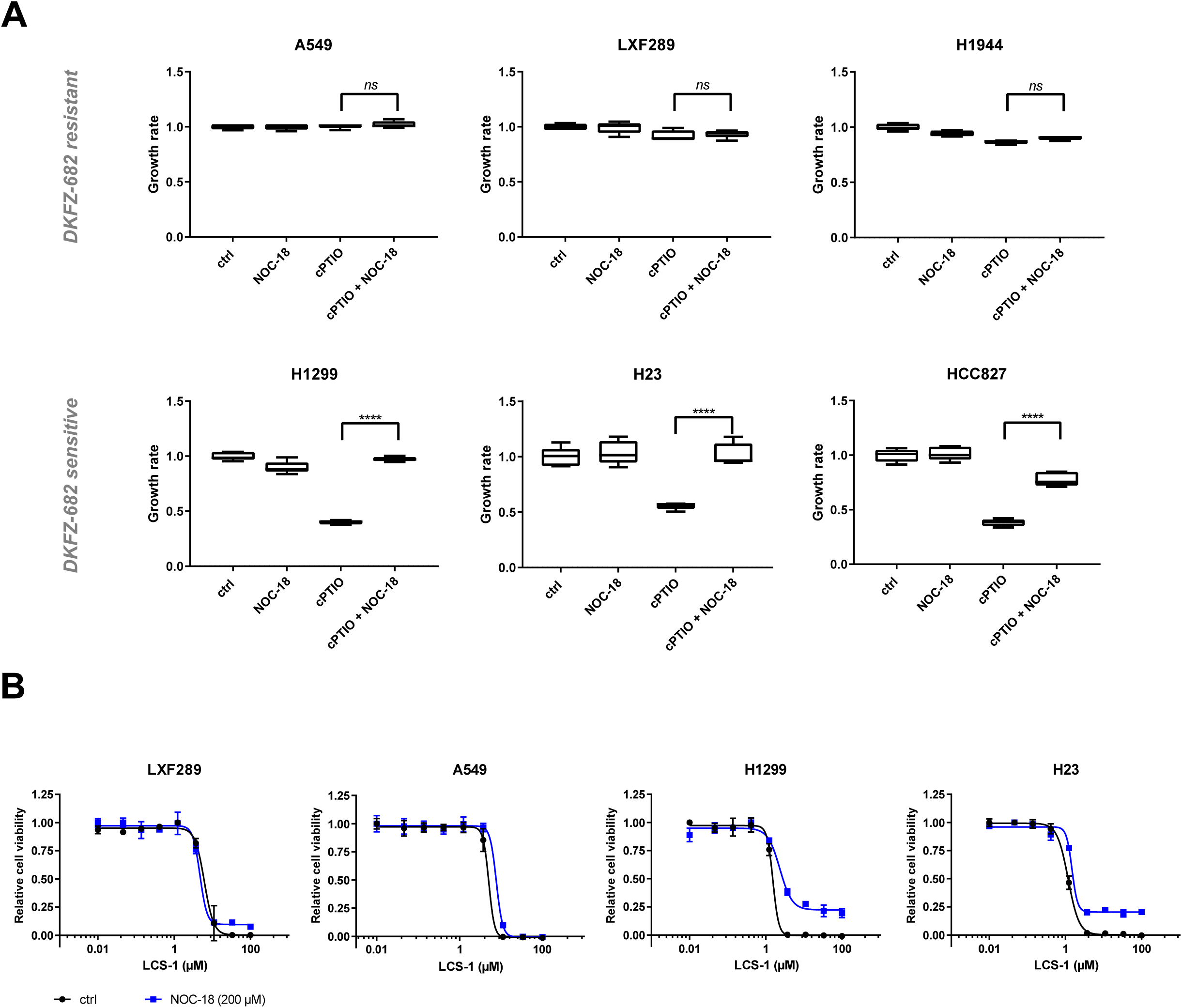
Nitric oxide (NO) depletion results in growth inhibition and superoxide susceptibility in cells with high RNS. **(A)** A549, LXF289, H1944, H1299, H23 and HCC827 cells were treated with NO scavenger cPTIO (50 µM), NO donor NOC-18 (40 µM), or a combination of both for 72 h. Growth rates were calculated based on nuclei count and presented as box plots (\*\*\*\**p*<0.0001, *ns*, not significant, two-tailed unpaired *t* test using the original data). The results are representative of three independent experiments**. (B)** LXF289, A549, H1299 and H23 cells were treated with a series of concentrations of SOD1 inhibitor LCS-1 for 24 h in the presence of NOC-18 (200 µM). Cell viability was quantified by the CellTiter-Blue assay. Bars represent results from three technical replicates. The results are representative of two independent experiments.

In addition to its role in cell signaling, it has previously been speculated that NO acts as a free radical scavenger, by reacting with O_2_^•-^ [30]. We tested this notion by blocking SOD1 with LCS-1 [36] in order to increase O_2_^•-^ to toxic levels. LCS-1 toxicity could be partially rescued by NOC-18 in sensitive cells (H1299, H23), supporting the notion that high RNS are a means to maintain ROS homeostasis favorable for cell proliferation (Fig. 3B). The cytoprotective role of NO was however insufficient to confer resistance against DKFZ-682 (Supplementary Fig. S3), suggesting that elevated NO levels function in ROS homeostasis rather than ROS defense.

### Sensitive cells have low expression levels of antioxidant capacity genes

Traditional assessment of target dependency for the development of redox drugs assumes that high expression levels of a target or pathway indicate a high dependency and vulnerability to inhibition. To identify biomarkers that correlate with sensitivity to ROS-inducing drugs in the NSCLC panel, we performed association analysis between published RNAseq, proteomics, metabolomics, and DNA-methylomics data [DepMap] and the EC50 values of DKFZ-682, determined for our cell line panel. We determined *p*-values for gene-by-gene Pearson correlation between expression or protein levels with EC50, and combined the ranks of both *p*-values by taking their average. Using a stringent cut-off (<38) on the average rank, we found that drug sensitivity is marked by low expression of proteins involved in NADP-regeneration (ME1, PGD, UGDH), as well as in antioxidation (GCLM, GSR, SLC7A11, TXN, AIFM2) and detoxification processes (CBR1, BLVRB, AKR1C1, AKR1C3, PTGR1, ALDH3A1, CYP4F11) (Fig. 4A). We named this group of 15 genes the <u>A</u>ntioxidant <u>C</u>apacity <u>B</u>iomarker (ACB) as they constitute the entire pathway involved in redox homeostasis and ROS stress defense. Unexpectedly, TXNRD1, the target of DKFZ-682, was not within the top 50 genes correlating with drug sensitivity. Average ACB expression and sensitivity to DKFZ-682 correlated with r=0.86, *p*≤0.0001 and was therefore highly predictive for drug response in NSCLC cell lines. The correlations were validated by our own RNAseq data [37] (r=0.79),with validated antibodies for a selection of ACB proteins (r=0.80, Supplementary Fig. S4) and MS-analysis (r=0.83), confirming the robustness of our findings. The fact that ACB members are occupying most of the critical nodes of ROS scavenging and detoxification pathways of reactive intermediates suggests that the highly coordinated expression pattern reflects a functional, interdependent network and that drug resistance cannot be induced by the upregulation of individual ACB members. We tested this by overexpression of single ACB genes such as *GSR*, *PTGR1*, *CBR1*, *TXN*, *GCLM* or *UGDH* in sensitive cells and observed no increase in resistance to DKFZ-682 (Supplementary Fig. S5A, S5B).

**Figure 4.**
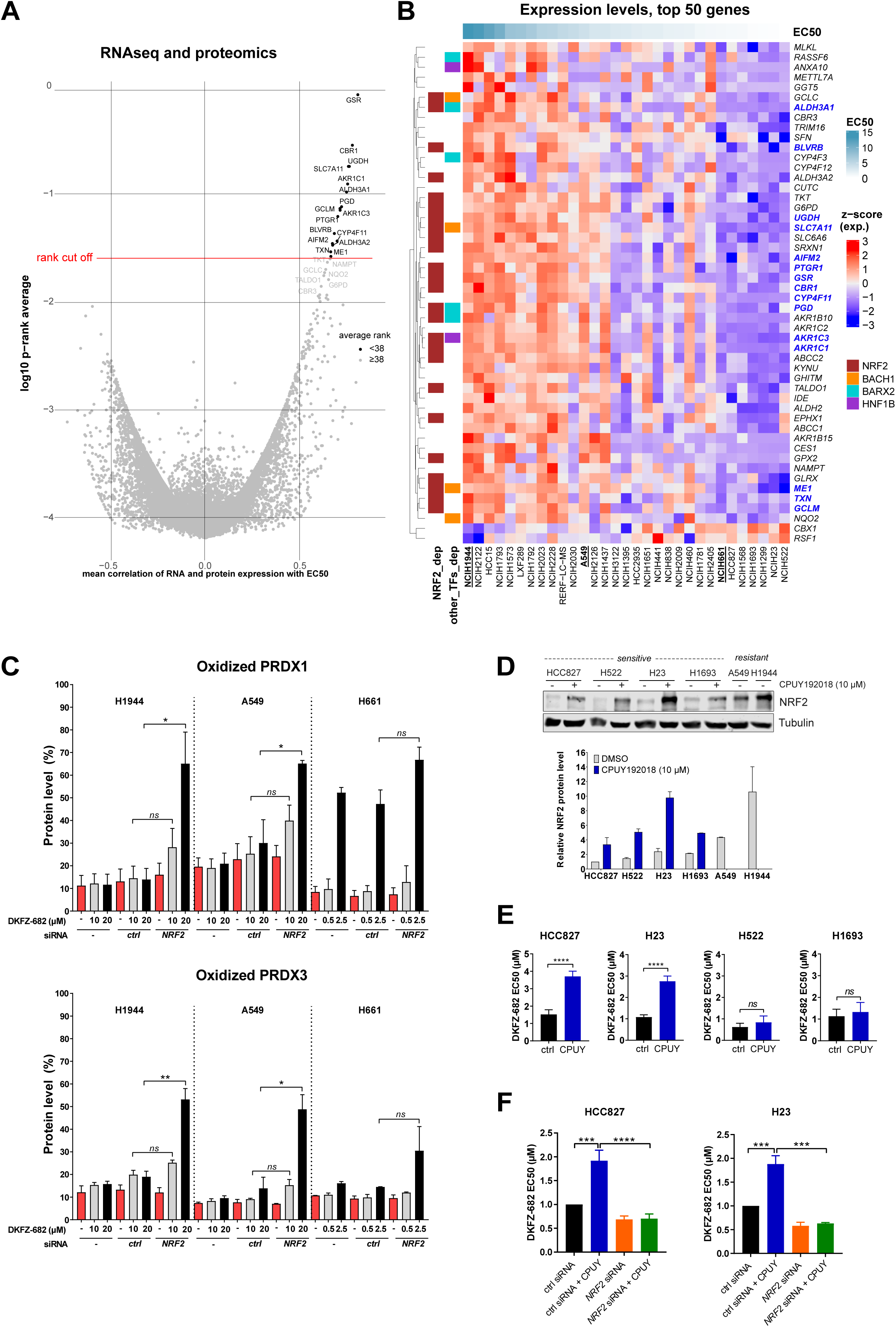
Identification of genes responsible for resistance to DKFZ-682. **(A)** Volcano plot of combined correlation of EC50 values with transcriptome and protein data of NSCLC cell lines. For each gene in overlap between expression and proteomics datasets, downloaded from DepMap project, we calculated average Pearson correlation coefficient and average ranks of *p*-value and plotted them. **(B)** Heatmap with the mRNA expression values of 50 genes that have the lowest average ranks from the correlation test. Expression values are scaled (z-score). Rows (genes) are clustered using “complete linkage” method and “Euclidean” distance. Row annotation shows some transcription factors that are common for these genes, as calculated via Dorothea R package. **(C)** H1944, A549 and H661 cell lines were transfected with nonsense (*ctrl*) or *NRF2* siRNA for 48 h and then treated with the indicated concentration of DKFZ-682 for 3 h. Oxidized (ox) and reduced (red) levels of PRDX1 and PRDX3 proteins were analysed by immunoblotting. Bar diagrams summarize the quantitative results from independent experiments (n=2-3, error bars indicate SD; \**q*<0.05, \*\**q*<0.01, *ns*, not significant, two-tailed unpaired *t* test). **(D)** Cell lines were treated with CPUY192018 (10 µM) or DMSO for 24 h. NRF2 protein level was analyzed by immunoblotting. Protein expression in DMSO-treated HCC827 was set to 1. Relative data represent mean of independent experiments (n=2, error bars indicate SEM). **(E)** Cells were seeded in 96-well plates and pre-treated with DMSO (ctrl) or CPUY192018 (CPUY, 10 µM) for 24 h. Then the cells were treated with a concentration series of DKFZ-682 for 24 h and the cell viability was measured by the CellTiter-Blue assay. **(F)** Cells were either treated with nonsense (ctrl) siRNA or siRNA against *NRF2*. The next day, cells were trypsinized and 7,500 cells were seeded into the wells of a 96-well plate. After 24 h, cells were treated with CPUY192018 (10 µM). After additional 24 h, cells were treated with a range of concentrations of DKFZ-682 for 24 h. Cell viability was assessed using CellTiter-Blue assay. Bar diagrams show the mean of EC50 data from independent experiments (n=4-8 **(E)**, n=3-4 **(F)**) each performed in triplicates (error bars indicate SD, \*\*\**q*<0.001, \*\*\*\**q*<0.0001, *ns*, not significant, two-tailed unpaired *t* test).

Taken together, these results demonstrate that sensitivity to ROS-inducing drugs is not conferred by the overexpression of the targeted redox-protein but by low expression levels of antioxidant capacity genes.

### Specific NRF2 levels are insufficient to establish low ACB status and drug sensitivity

Transcription factor mapping and knockdown experiments showed that the ACBs are regulated by NRF2 (Fig. 4B and Supplementary Fig. S6A). Consistent with the role of ACBs in ROS defense, *NRF2* knockdown significantly increased DKFZ-682-induced PRDX1/3 oxidation and drug sensitivity in resistant cells but not in the sensitive reference cell line H661 (Fig. 4C, black bars; Supplementary Fig. S6B, S6C). Surprisingly and contrary to the notion that high levels of NRF2 and its target proteins are required to buffer excessive steady state ROS production, the loss of NRF2 did not affect, baseline redox homeostasis in resistant cells (Fig. 4C, red bars).

Despite its prominent role as a master regulator of redox homeostasis and ROS stress defense [17], NRF2 protein levels showed a lower correlation with drug sensitivity (r=0.61, *p*=0.0002), as compared to ACB RNA or protein (r=0.86 or r=0.80, *p*<0.0001). This was particularly obvious in the H838 cell line, which demonstrated comparable sensitivity and low ACB expression similar to H441 and H2009, but expressed >5 times higher NRF2 levels. Inversely, H1792, which is among the highest ACB expressing and one of the most resistant cell lines in our panel, expresses NRF2 levels comparable to some sensitive cell lines (Supplementary Fig. S4). These discrepancies raise the question whether the ACB status in sensitive cells is not predominantly determined by NRF2 activity but rather by alternative mechanisms ensuring low ACB expression.

To answer this question, we first established whether sensitive cells have a functional NRF2-system. NRF2 protein and activity levels can be increased by CPUY192018 (CPUY), an inhibitor of the KEAP1-NRF2 protein-protein interaction [38]. Induced NRF2 levels were comparable to those detected in the most resistant cell lines (Fig. 4D) and were functional in sensitive cells since they resulted in the induction of *SRXN1*, a commonly used proxy for NRF2 activity [39] (Supplementary Fig. S7A). Surprisingly, despite a strong NRF2 protein induction, we observed only a moderate decrease of drug sensitivity in some cells (H23 and HCC827) while others remained sensitive (H522 and H1693, Fig. 4E). Consistent with this, NRF2 induction did not alter the redox buffer capacity of H522 and led to a moderate increase only in HCC827 and H23 (Supplementary Fig. S7B). The CPUY effect in the responder cells was dependent on NRF2 protein as a knockdown of *NRF2* prevented the induction of resistance (Fig. 4F). Global expression analysis of CPUY-treated cells (Supplementary Fig. S7C) revealed that in the CPUY non-responding cell line H522 all but one ACB (*GCLM*) remained silent while HCC827 demonstrated induction of four members of the ACB set (*AKR1C1*, *AKR1C3*, *GCLM*, *SLC7A11*) plus additional genes involved in redox homeostasis (GCLC and GPX2). Interestingly, although CPUY treatment induced a ∼10-fold lower number of genes by > 1.5-fold in H522, as compared to HCC827 cells, it led to a strong induction of *HMOX1* and *OSGIN1*, two reliable proxies for NRF2 activity [39]. The data presented so far are in line with our initial hypothesis that low ACB status and phenotype are not only established by low NRF2 levels, but also by a silencing mechanism for at least a subset of ACBs, possibly through the lack of positive transcription factors, the expression of transcription repressors or by epigenetic repression.

To elucidate the underlying mechanism, we first asked whether the low expression of ACBs could be explained by promoter methylation. Neither *in silico* analysis, using a stringent *p*-value cut-off (<0.001) for the DNA methylation vector (Supplementary Fig. S8A) nor pretreatment of sensitive cells with the DNMT1 inhibitor azacytidine (Supplementary Fig. S8B) hinted at a prominent role in ACB repression. Actionable histone-based mechanisms like deacetylation or methylation were also evaluated by pretreatment of cells with Class-1 histone deacetylase (HDAC) inhibitors PAOA, BRD-6929 or BML-210 as well as with histone N-methyltransferase (HMT) inhibitors BIX-01294 and UNC-0642 (Supplementary Fig. S8C). None of these enhanced CPUY induced resistance, suggesting that direct or indirect epigenetic suppression of ACB promoters does not contribute to the low-ACB status of sensitive cells.

Taken together, the low ACB status in sensitive cells is not primarily established by a lack of NRF2 or epigenetic regulation but by additional mechanisms leading to robust silencing of ACB genes.

### STAT3/5 contributes to low ACB expression and drug sensitivity

An alternative mechanism for low ACB expression could be the lack of positive transcription factors. We used gene set enrichment analysis of transcription factor target genes in CPUY responding versus non-responding cells and found that the NFKB subunits RELA, NFkB1 and REL might be involved in NRF2 dependent drug resistance (Supplementary Fig. S9A).

Contrary to our expectation, we did not blunt CPUY induced drug resistance of HCC827 when inhibiting NFkB activity with SC75741 [40] nor BAY11-7082 [41], arguing against a prominent involvement of NFkB in ACB expression (Supplementary Fig. S9B). Surprisingly, when using TPCA-1, which inhibits the phosphorylation and activation of both NFKB and STAT1/3 [42] (Supplementary Fig. S10A), we observed an enhancement rather than a reduction of drug resistance in HCC827, but not in H522 (Fig. 5A). This was reproduced by the pan-JAK inhibitor pyridone 6 (P6) [43] and a competitive STAT1/3 inhibitor C188-9 [44] (Supplementary Fig. S10A-C). Inhibition of STATs resulted in elevated ROS buffer capacity in an additive fashion with CPUY (Supplementary Fig. S10D). Except for C188-9, none of the inhibitors induced NRF2, arguing against a nonspecific induction of ROS (Supplementary Fig. S10A). Comparable to chemical inhibition, siRNA-induced knockdown of *STAT1*, *STAT3* or *STAT5A/B*, combined with NRF2 induction, caused a drop of drug sensitivity with *STAT5A/B* demonstrating even an additive effect (Fig. 5B). Using the corresponding cell line pair HCC827 and H522, we found that STAT inhibition by C188-9 enhanced the expression of 5 ACB members beyond the induction achieved by CPUY treatment in HCC827 but only 2 ACB members in H522 (Fig. 5C). The repressive activity of STATs was observed not only in the EGFR-mutant HCC827 [45], but also to some extent in H23, H661 and H1299, expressing wt EGFR (Supplementary Fig. S10E).

**Figure 5.**
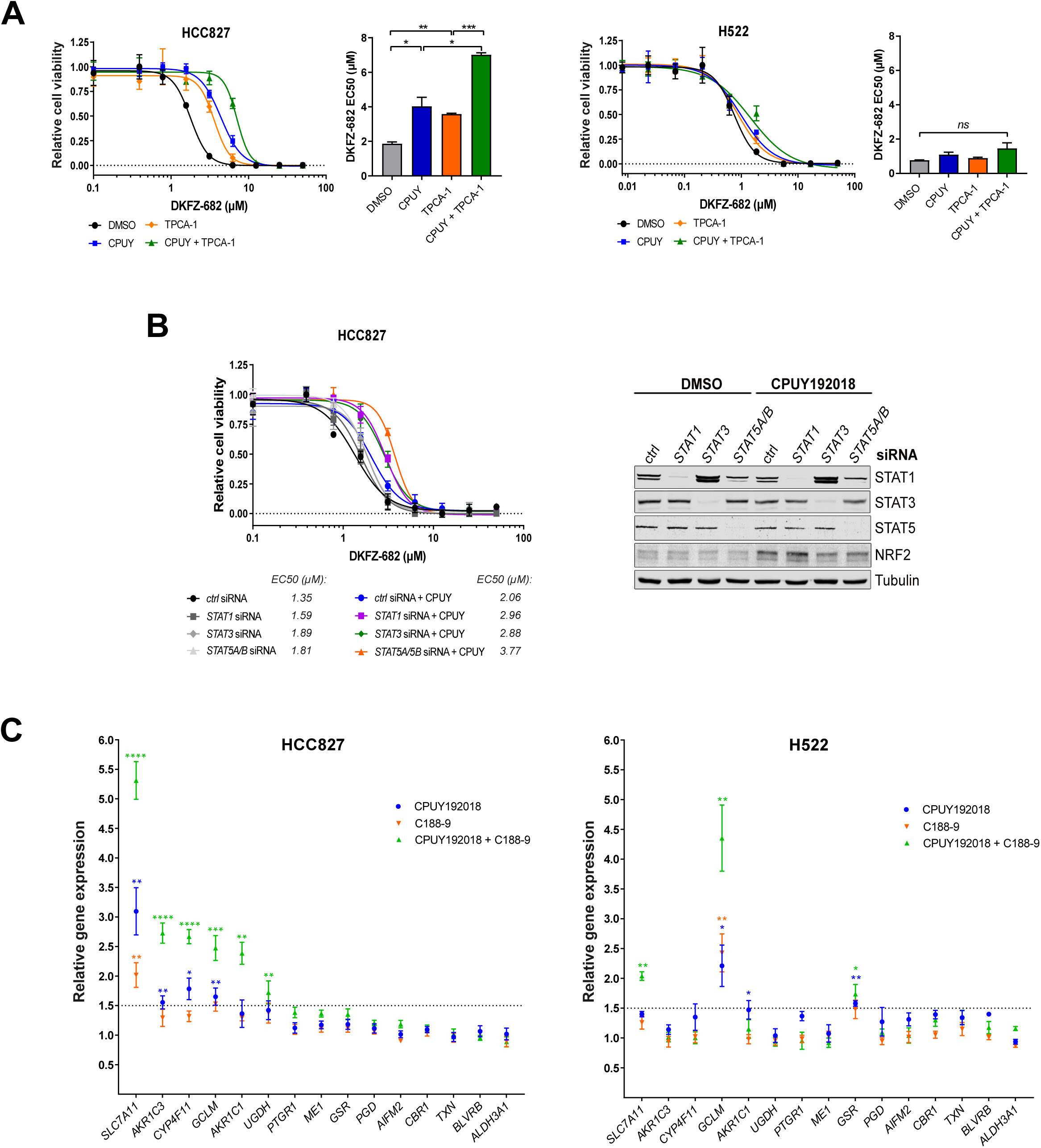
Inactivation of STAT enhances cell resistance to ROS inducing drug. **(A)** HCC827 and H522 cells were treated with 10 µM CPUY192018 (CPUY) and 1 µM TPCA-1 or a combination of both for 24 h. Next, the cells were treated with series dilutions of DKFZ-682, and after 24 h, cell viability was quantified by the CellTiter-Blue assay. Bar diagrams show the mean of EC50 data from independent experiments (n=2) each performed in triplicates (error bars indicate SD, \**q*<0.05, \*\**q*<0.01, \*\*\**q*<0.001, *ns*, not significant, two-tailed unpaired *t* test). **(B)** HCC827 cells were transfected with siRNA directed against *STAT1* (10 nM), *STAT3* (10 nM), *STAT5A/B* (5 nM + 5 nM) or nonsense (ctrl, 10 nM) siRNA overnight and then seeded in 96-well plates (7,000 cells/well) and pretreated with DMSO or CPUY192018 (CPUY, 10 µM) for 24 h. Then the cells were treated with a concentration series of DKFZ-682 for 24 h and the cell viability was measured by the CellTiter-Blue assay. The graph is representative of two independent experiments each performed in triplicates. STAT protein expression was analysed by immunoblotting. Representative western blots are shown. **(C)** RNAseq analysis shows the expression of ACB genes in the HCC827 and H522 cells treated with CPUY192018 (10 µM) and/or C188-9 (10 µM) for 6 h. Results are shown as fold change compared with DMSO treated control. Results are representative of two independent experiments each performed in triplicates (error bars indicate SD, \**q*<0.05, \*\**q*<0.01, \*\*\**q*<0.001, \*\*\*\**q*<0.0001, unpaired multiple *t* test, comparison DMSO versus drug treatment).

Our results show that sensitivity against ROS-inducing drugs is not only due to low NRF2 levels, but also mediated by the repressive activity of STAT proteins.

### ACB biomarkers stratify drugs to sensitive cells

An essential step in assessing the potential of the ACB set to match a susceptible patient with an appropriate drug is to correlate the biomarker expression with a treatment outcome. Currently there are no published expression data from NSCLC patients or PDX models that have been treated systematically with ROS-inducing drugs. Therefore, we used an indirect way to validate the predictive power of the ACB set by asking whether the biomarker profile, which has been identified through the selective TXNRD1 inhibitor DKFZ-682, is able to identify ROS-inducing drugs in a large collection of tool compounds and clinical drugs tested in NSCLC adenocarcinoma cell lines. To this end, we used data from the CTRP project [19] and calculated correlations between the expression of each gene of the ACB set and the area under the curve-drug response values (AUC). The resulting 15-element vectors of each of the 543 compounds and of DKFZ-682 were clustered to obtain the Euclidean distance of each drug to DKFZ-682 as a measure of similarity (Fig. 6A). ACB exhibited a high predictive power since we obtained an enrichment of compounds reported to induce ferroptosis (ML210, ML162, 1S,3R-RSL 3, erastin), bind to TXN (necrosulfonamide, PX-12, PRIMA-1), or inhibit TXNRD1 (piperlongumine and WP1130). We validated the predicted activity profile of selected compounds and obtained a high correlation of drug responses with the activity profile of DKFZ-682. Furthermore, we could confirm that most of the identified compounds induced ROS in sensitive cells (Supplementary Table S3).

**Figure 6.**
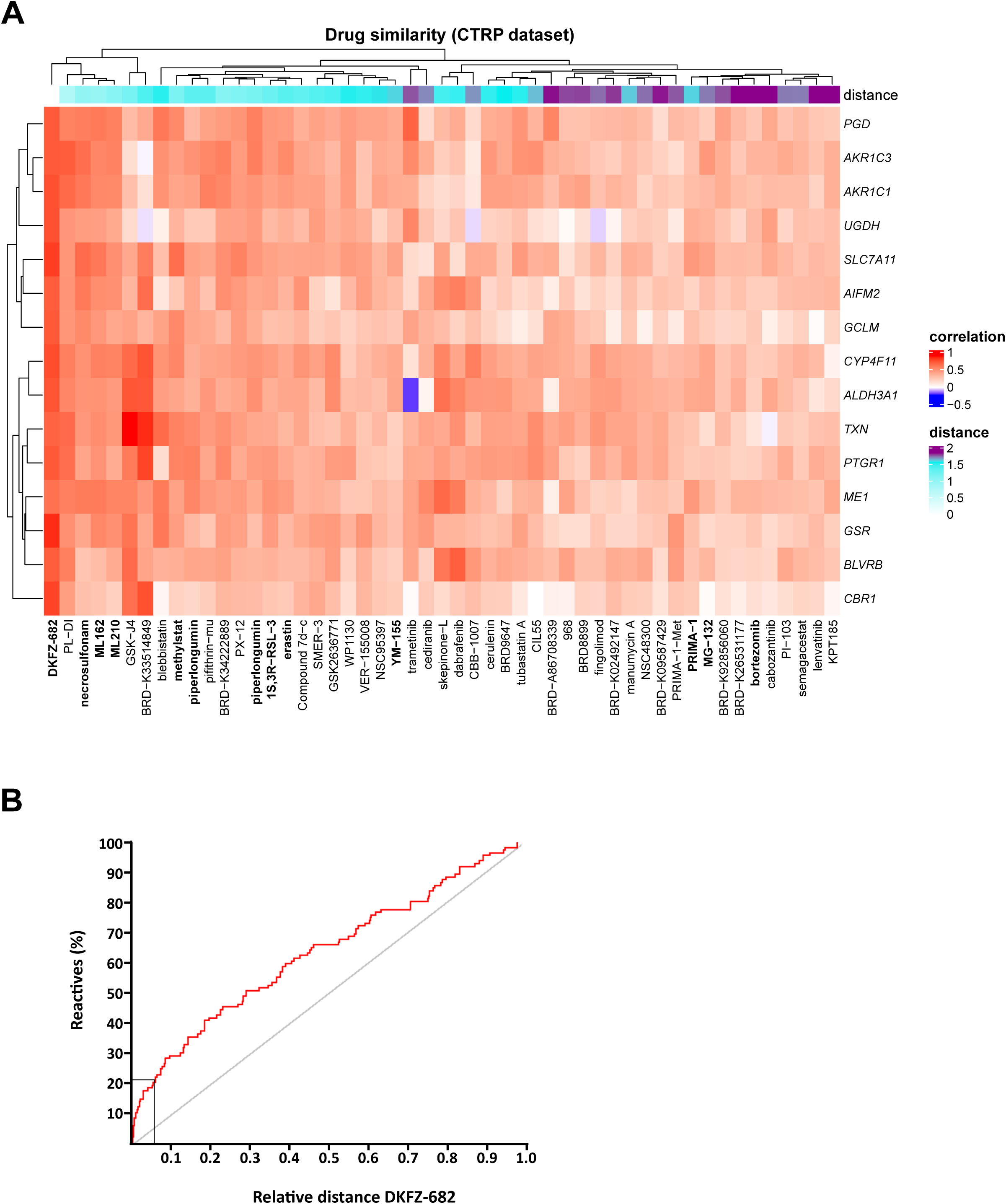
The correlation of ACB and drug activity profiles reveals ROS inducing drugs. **(A)** We calculated correlations between AUC (area under the curve) and expression of ACB genes for all compounds in CTRP dataset in 61 NSCLC adenocarcinoma cell lines. Then having correlation values for DKFZ-682 (correlation between EC50 and expression values), we calculated Euclidean distance between DKFZ-682 and all CTRP compound in “ACB correlation” space. The heatmap shows correlation values between AUC and ACB expression values for 50 compounds with the shortest distance from DKFZ-682. Columns (compounds) and rows (genes) are clustered using “complete linkage” method and “Euclidean” distance. The distance from DKFZ-682 is shown as top annotation. **(B)** The ROC curve (Receiver-Operating-Characteristics) shows the enrichment of reactive groups in compounds with shorter distances to DKFZ-682. All 543 substances were classified either to have a chemical reactive group (125 compounds, e. g. Michael acceptor) or to be chemically inconspicuous. Up to a Euclidean distance < 2 (relative distance to DKFZ-682 of 0.06, 33 compounds), we identified 21 reactive compounds (indicated by rectangle). Compared to just randomly selecting compounds this is an enrichment of almost three, supporting the idea that reactivity of a compound can lead to the described drug activity profile. The ROC curve was generated showing on the Y-axis the percentage of identified compounds with reactive features. The X-axis displays the sorted distances.

A closer inspection of the chemical structures revealed that most of the top 50 drugs contain reactive pharmacophores or functional groups, such as Michael acceptors or alpha-chloro amides, which are likely to react with nucleophiles, in particular cysteines or selenocysteines. We verified this by testing for the enrichment of reactive groups in compounds with a shorter Euclidean distance to DKFZ-682. The results confirm that reactive, rather than randomly selected, compounds tend to have a drug activity profile comparable to DKFZ-682 (Fig. 6B).

Interestingly, both ferroptosis defense genes [46] (ACB members *GSR*, *AIFM2*, *TXN*, *GCLM*, *AKR1C3* and *SLC7A11*) and ferroptosis-inducing drugs were identified in this study. This raised the question whether a distinct pattern of polyunsaturated fatty acids (PUFA) [47], e. g. PUFA^high^ versus PUFA^low^, would correlate with the sensitivity to drug-induced ROS and could serve as an additional biomarker. However, correlation analysis of previously published data [15] with our EC50 data showed, that drug sensitivity was independent of lipid composition (Supplementary Fig. S11A). Consistent with this, sensitive cells did not demonstrate enhanced lipid peroxidation upon drug-induced ROS nor did they show higher free iron levels than their resistant counterparts (Supplementary Fig. S11B, S11C). Testing of IKE (carbonyl erastin analog), a more drug-like derivate of erastin [48] in a panel of 18 NSCLC cell lines demonstrated a more than 500-fold selectivity for low-ACB, compared to high-ACB cells. Interestingly, the entire ACB set demonstrated a better correlation with IKE activity than the set of ferroptosis defense genes or AIFM2 alone (Supplementary Table S4). These data suggest that the development of this compound class would greatly benefit from patient stratification and that the ACB set could be instrumental in this.

### ACBs predict drug response in patient-derived tumors

In order to evaluate the relevance of ACBs beyond cell lines, we first evaluated whether ACB genes are expressed in a coordinated fashion not only in our cell line panel but also in tumor biopsies (RNAseq data derived from the TCGA-LUAD cohort and proteomics data from Lehtiö et al. [49]), or a collection of 59 PDX models from NSCLC patients [50]. A gene-by-gene correlation of ACB expression revealed that the majority of biomarkers were similarly expressed in LUAD tumors as in PDX models and the cell lines, indicating that the underlying control of ACB expression is robust across models and patient tumors (Fig. 7A; Supplementary Fig. S12). ACB expression in normal lung tissue was less coordinated, suggesting that LUAD has acquired this characteristic expression profile during tumorigenesis.

**Figure 7.**
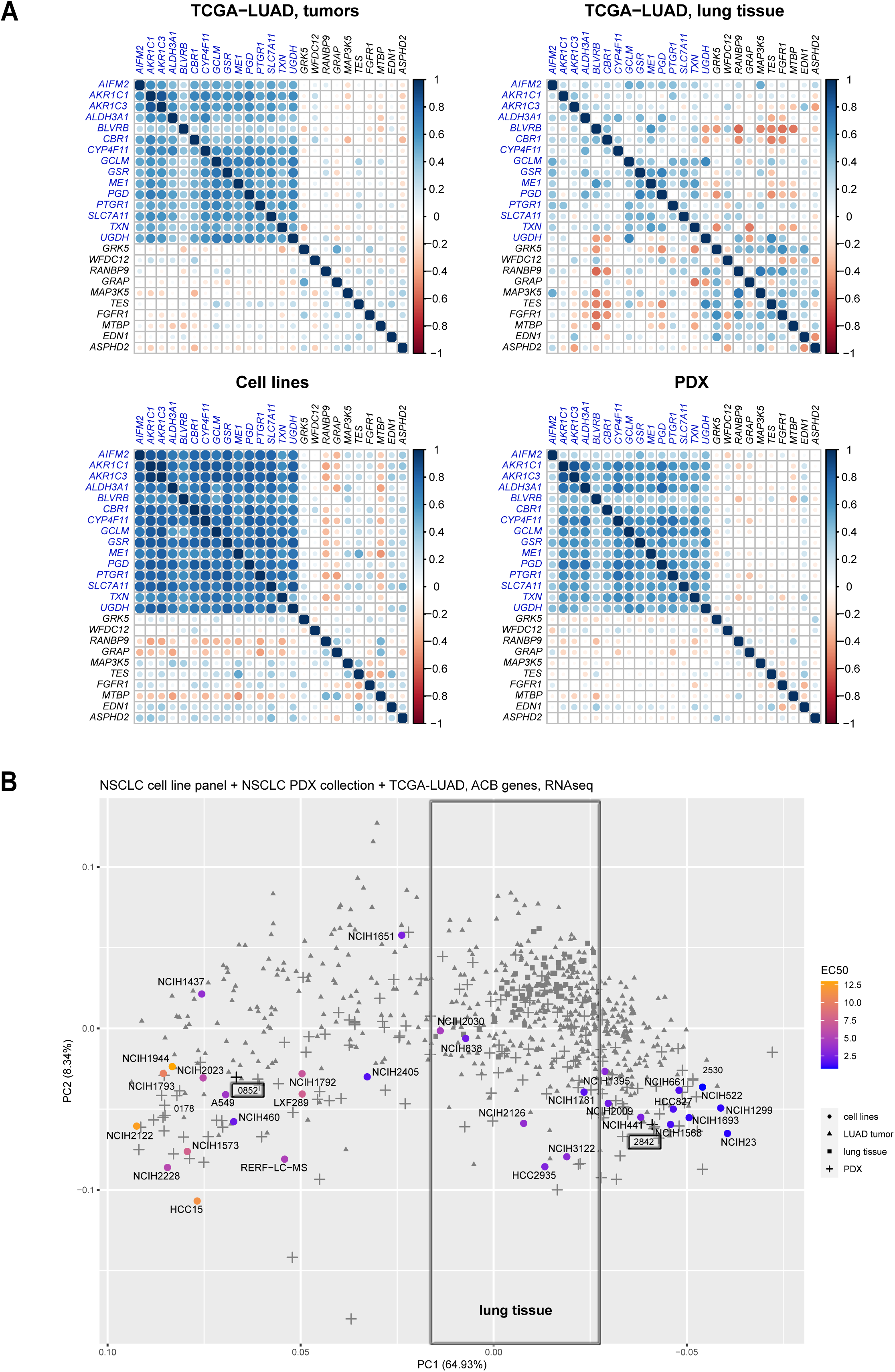
Coordination and expression space indicates conserved function in cell lines and tumors. **(A)** Correlation plots show gene-gene correlations for ACB genes in 576 TCGA-LUAD tumors, TCGA-LUAD lung tissue, 31 NSCLC cell lines and 59 PDX models. A group of non-ACB genes is included for comparison. **(B)** Principal component analysis plot (1^st^ and 2^nd^ PCA components) on ACB genes. Samples include NSCLC cell lines, PDX models, TCGA-LUAD tumors and TCGA-LUAD lung tissue samples. Sensitivity of cell lines is indicated by the color code (blue – sensitive, orange – resistant).

To test the predictive power of the ACB set for DKFZ-682, we used *ex vivo*, drug sensitivity testing in patient derived tumor material. A principal component analysis (PCA) revealed that PDX tumors exhibit a wide span of ACB expression, comparable to the cell line panel (Supplementary Fig. S13A). We selected a low-ACB model (CTG-2842) and a tumor model clustering with high ACB cell lines (CTG-0852). *Ex vivo* PDX drug sensitivity assay showed that the low-ACB model was indeed more sensitive to drug treatment (Supplementary Fig. S13B). It should be noted that the drug effect range of PDX models was considerably lower and that the variation among the biological repeats was higher than we observed under highly standardized conditions in the cell line panel. Most likely, this was caused by the infiltration of PDX tumor tissue with host cells and by a larger size variation of 3D cell aggregates in the test samples. An additional confirmation of the predictive power of the ACB set has recently been provided by Yan et al. In their in *vivo* PDX models, low levels of GSR and coexpressed antioxidant genes correlate with higher sensitivity to ROS-inducing drug auranofin [51].

### ACBs allow to identify patient tumors with low ROS buffer capacity

In order to translate from in vitro models to cancers, we then asked whether ACBs populate the expression space charted by cell lines and PDX samples also in patient tumors. PCA analysis, using the ACB expression of the combined data sets, revealed that normal lung tissue showed a distinct expression profile of ACBs, suggesting that selected, low-ACB patients could benefit from a usable therapeutic window for the treatment with ROS-inducing drugs (Fig. 7B). A hierarchical cluster analysis revealed that less than 1 % of the 576 patients in the TCGA-LUAD cohort demonstrated ACB expression comparable to the most sensitive cell lines and PDX models (Supplementary Fig. S13C), underscoring that a clinical trial in LUAD, without patient stratification, is unlikely to demonstrate any clinical benefit of ROS-inducing drugs.

In contrast to LUAD, ACBs are favorably expressed in a high proportion of patients with acute myeloid leukemia (LAML), uveal melanoma (UVM), diffuse large b-cell lymphoma (DLBC), and pheochromocytoma and paraganglioma (PCPG), suggesting that more high responders to ROS-inducing drugs can be identified in these entities when ACB-stratified in clinical trials (Fig. 8A). We validated this hypothesis with the example of LAML. Conveniently, the space of ACB expression in AML cell lines characterized in the CCLE project is comparable to the TCGA-LAML cohort, which allowed us to assemble a representative panel of 17 cell lines to determine the activity profile of DKFZ-682 (Supplementary Fig. S14). As predicted, the AML cell panel, which expresses substantially lower levels of ACBs, demonstrates significantly higher average drug sensitivity than the NSCLC panel (Fig. 8B). The correlation of drug sensitivity and 15 ACB genes expression within the AML cell panel was r=0.55 (*p*=0.003), arguing for a functional role of the ACB set in drug sensitivity within this susceptible entity. Using an iterative approach to eliminate ACB components with minor contribution to drug sensitivity correlation (*AKR1C1*, *BLVRB*, *GSR* and *PTGR1*), we observed a correlation of r=0.69 (*p*=0.002) between expression of the remaining 11 ACB genes and sensitivity of AML cells to DKFZ-682. It is worth noting that the entire set of ACBs, calculated as average ACB expression, demonstrated a far stronger correlation than any of the individual components, underscoring our notion that combining multiple, functionally connected markers mitigates the risk associated with variable expression of individual markers.

**Figure 8.**
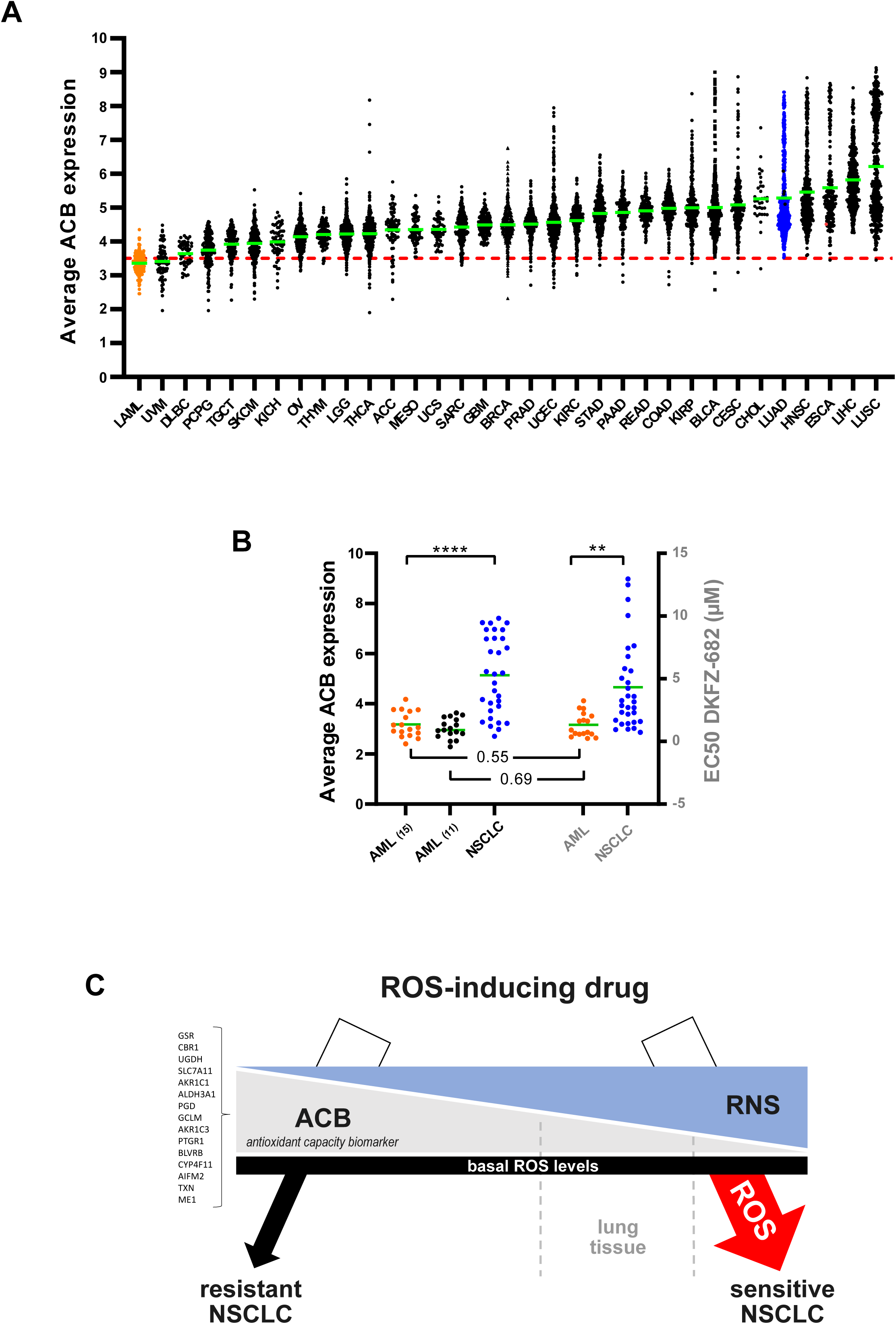
ACB expression levels across tumor entities reveal substantial heterogeneity in ROS buffer capacity and suggest cancers with favorable ACB profiles. (A) Inferred ROS buffer capacity based on the average expression of ACBs (antioxidant capacity biomarker) across >11,000 tumors profiled by The Cancer Genome Atlas. Each dot represents the average expression of the ACB set of 15 genes in an individual tumor. The red line represents the ACB level of H23, one of the most sensitive cell lines in our NSCLC cell panel. Green lines represent the mean value for each cancer type. The cancer entity LUAD, corresponding to our panel of lung adenocarcinoma cell lines, is marked in blue, LAML tumors, corresponding to our validation panel of AML cell lines are marked in orange. Y-axis shows the average expression of the ACB set. TCGA expression data were converted from RPKM to log_2_(TPM+1) units to match the value of H23. **(B)** Comparison of average ACB expression and drug sensitivity (EC50) in the cell line panels representing NSCLC adenocarcinoma (blue), AML_(15)_ (15 ACB genes; orange) and AML_(11)_ (11 of 15 selected ACB genes; black). Highly significant differences in ACB expression (**** *p*<0.0001, two-tailed unpaired *t* test) and EC50 values (** *p*=0.0012, two-tailed unpaired *t* test) are indicated. The Pearson correlation of expression of the original set of 15 ACBs and drug sensitivity within the AML panel is r=0.55, *p*=0.003. The correlation of the reduced set of 11 ACBs (without AKR1C1, BLVRB, GSR and PTGR1, see Supplementary Table 10) with drug sensitivity is r=0.69, p=0.002. **(C) Biomarker profile of cancer cells responsive to redox-targeting drugs**. Cancer cells sensitive to ROS-inducing drugs demonstrate high levels of RNS and low expression levels of ACBs. Tumor cells have comparable basal ROS levels. Upon drug insult, resistant tumor cells and most normal tissues are able to buffer ROS induction to prevent cell damage and apoptosis. Sensitive cells, due to their low ROS-buffer capacity, experience a deadly level of ROS-damage. The susceptibility of normal lung tissue to ROS-inducing drugs can be expected to be variable and dependent on pharmacological properties of the substance.

Taken together, we have identified tumor entities with predominantly low ACB expression and show that this correlates with high sensitivity to DKFZ-682.

## Discussion

Predictive biomarkers are instrumental in the success of targeted cancer therapies. They help to identify patients whose tumors have the right genetic background to respond more effectively to the drug than normal tissues. The lackluster success of drugs targeting the ROS-scavenging machinery in clinical trials suggests that differential ROS sensitivity might be absent in most cancers and that precise identification of ROS-sensitive tumor entities or a selection of patients within a given cohort would be instrumental for the clinical development of ROS-inducing drugs. Currently, there are no published omics data from patients or PDX cohorts of any cancer entity that have been treated systematically with ROS-inducing drugs and that could be used to directly identify efficacy biomarkers.

Our study’s approach to identify biomarkers for hypersensitivity to ROS-inducing drugs was based on components that enable high-resolution observations. First, we chose DKFZ-682 rather than the less selective auranofin to reduce phenotypic noise resulting from its high nonspecific reactivity with exposed cysteines in proteins and glutathione and its limited selectivity for the TXN system [27, 52]. Second, we used NSCLC cell lines to assemble a cell panel that spanned a wide and continuous range of drug sensitivities and provided in-depth omics data suitable for correlation analysis. Using combined RNA/protein data, rather than only one parameter, plus a very stringent rank cut-off allowed us to identify a set of biomarkers with an unprecedented high correlation to drug sensitivity.

Predictive biomarkers should fulfill two main criteria. Firstly, they should be easily detectable and quantifiable. Secondly, they should have a clear rationale in that they are tightly coupled with the specific mechanism of drug action [53]. Regarding the first criterion, using the ACB set would require state-of-the-art RNA quantification of patient tumor biopsies. A similar approach is currently being applied to a set of 10 RNA-based drug response predictor genes (DRP) [54] which is now used to identify patients likely to benefit from treatment with the alkylating drug irofulven [55]. When validating the biomarker concept in a non-NSCLC entity, we could confirm that the ACB set correctly predicts high drug sensitivity in AML cells. A reduction of the ACB set of 15 genes to 11 genes substantially improved the correlation of drug sensitivity and ACB expression within the cell line panel. For future drug development projects, we expect that more detailed studies will be required, especially in patient-derived cells and tumor models, to fine-tune entity-specific ACB sets.

As for the second criterion (a clear rationale), our findings show that hypersensitive NSCLC cells are characterized by low expression levels of most components required for defense against induced ROS stress. Those genes can be structured into four functional categories: (1) proteins involved in the synthesis of glutathione (GSH); (2) enzymes regenerating NADPH and GSH; (3) genuine ROS scavenging proteins, which use NADPH or GSH to transfer electrons to ROS, cysteine-oxidized proteins, and lipid radicals; (4) enzymes involved in xenobiotics response and reduction of toxic intermediates. Individual members of ACBs have previously been discussed as indicative of sensitivity to ROS-inducing compounds in NSCLC cell lines, like GSR [51], AIFM2 [56], GCLM, and GCLC [57]. Our data demonstrate a highly coordinated expression of ACB genes, which suggests that the observations reported in these studies, may be based on the expression of the ACB set rather than on individual components.

Contrary to our expectation, TXNRD1, the target of DKFZ-682, was not within the top 50 genes correlating with drug sensitivity. This indicates that for ROS-inducing drugs, the steady state ROS defense capacity, composed of multiple components, may be a stronger determinant of drug efficacy than the expression level of the actual target. Our data also show that high expression levels of ACB are not a sign for a druggable dependency on ROS buffer capacity but rather an obstacle for treating such tumors with redox targeting drugs. In this sense, the redox system and individual targets need to be considered as pan-essential genes, whose therapeutic potential can be reduced by dose limiting toxicity in normal tissue or insufficient target engagement resulting from protein overexpression in tumor tissue [58]. Accordingly, we propose to abandon the traditional assessment of target dependency for the development of redox drugs, which assumes that high expression levels of a target or pathway indicate a high dependency and vulnerability to inhibition. Rather, drug sensitivity is a function of ROS buffer capacity, which, in the case of sensitive NSCLC cells, is marked by constitutive repression of ACBs and high levels of RNS (Fig. 8C).

We found that compounds inducing ROS via alternative targets, like ferroptosis inducers, are most active in cells expressing low levels of ACBs. This appears to also be the case in drug-tolerant persister cells, which demonstrate global repression of antioxidant genes [59]. Ferroptosis inducing strategies are attracting particular interest due to their detailed understanding of the underlying mechanisms and the druggability of key components of the pathway. However, despite the wealth of preclinical and clinical compounds inducing ferroptosis, toxicity and off-target effects remain a challenge in clinical oncology [60]. Based on our data, we suggest using ACBs to determine which tumor types are most suitable for ferroptosis-based therapies.

Our study shows that resistant and sensitive cells have established comparable redox homeostasis to support cancer-specific hallmarks. This finding raises the question of how sensitive cells manage to establish intracellular eustress [61], optimal for most cellular functions, without expressing appreciable levels of ROS-scavenging proteins. NO can act as a ROS scavenger [62] and thereby potentially compensates for low ACB expression. Although this has not been described before for ROS-sensitive cancer cells, the underlying chemistry clearly supports this notion in that NO acts as a free radical scavenger by reacting with O_2_^•-^, to form the non-radical, slow-reacting molecule ONOO^-^ [30]. NO was also reported to protect cells against induced ROS stress [63] and ferroptotic cell death [64]. Our data show that, although NO can reduce the burden on the enzymatic ROS-scavenging machinery in sensitive cells to some extent, it is insufficient to abrogate the cytotoxicity of DKFZ-682. Also, we observed that NO levels are reduced upon drug treatment and speculate that sensitive cells face a dual handicap in their defense against ROS. When TNXRD1 is inhibited, they lack the capacity to deal with the dramatically increased H_2_O_2_ levels, which, on the one hand, leads to the formation of toxic OH^•^ via iron and copper-mediated Fenton reaction. On the other hand, H_2_O_2_ can directly react with NO to form additional HO^•^ [34], increasing oxidative stress even more. In addition to directly supporting redox homeostasis in the context of a weak ROS-scavenging machinery, RNS might also act as a regulator of protein functions, through post-translational modifications of cysteine and tyrosine residues [65]. Interestingly, tyrosine nitrosation appears to increase the activity of the ROS scavenging components PRDX2 [66] and MGST1 [67] which have been reported to promote cell proliferation in NSCLC lines [68, 69].

Another important question raised in this study concerns the molecular mechanism involved in the low-ACB status of sensitive cells. Our data provide a first insight by showing that STAT3 and STAT5A/B activity represses ACBs and enhances drug sensitivity. Interestingly, a similar role of STAT proteins in ROS homeostasis has previously been reported in chronic myelogenous leukemia (CML) [70], breast cancer cell lines [71] and normal tissue [72]. However, our data also show that the repressor function of STATs applies only to a subset of ACBs and only to a part of sensitive cells and therefore fall short of providing a generalizable mechanism. More detailed studies to unravel the potentially complex mix of several mechanisms would offer the opportunity to identify actionable nodes which could allow inducing drug sensitivity in entities with a less favorable ACB status.

The ACB concept presented in this study can be used the estimate what percentage of patients in a given cancer entity are likely to respond to ROS targeting therapies. Based on the ACB levels of the most sensitive NSCLC cell lines, only a low single-digit percentage of patients of the TCGA_LUAD cohort would qualify as potential high responders. This hypothesis can be refined with the help of a recent proteomics study, which showed that tumors from large-cell neuroendocrine carcinoma (LCNEC) express the lowest levels of ACB proteins in the NSCLC entity [49]. However, the number of responder patients could be expanded by drug combinations; this follows the rationale that most drug treatments induce ROS in cancer cells [4], and would respond synergistically when their ROS scavenging capacity is inhibited or challenged in addition. Targeted therapies like tyrosine kinase inhibitors (TKI) increase oxidative stress to a level exhausting the reductive capacity of cancer cells [73], exemplified most clearly by studies on axitinib, sorafenib, erlotinib, vemurafenib and crizotinib [74]. The latter report is particularly instructive as it showed that combination with disulfiram strongly enhanced crizotinib efficacy. Disulfiram blocks the ROS-scavenging and detoxification mediated by ALDH isozymes [75]. In addition, it forms copper complexes *in vivo*, a reaction that results in the generation of ROS, making disulfiram a broad challenger of the ROS-scavenging machinery [76]. It is tempting to speculate that in patients with a low-ACB profile, a combination of disulfiram and TKI like crizotinib could be highly efficacious.

As stated earlier, the clinical development of dedicated, redox-targeting drugs has so far been plagued with disappointing success rates. Like in any other therapeutic class, the reasons can be manifold and need to be assessed on an individual basis; however, the majority of redox-targeting compounds contains electrophilic functional groups, and engage their targets via covalent bond formation. Assuming that physicochemical properties, which affect stability and distribution, are not an intrinsic liability of redox-targeting drugs and, therefore, can be improved in the course of lead optimization, the main hurdle remaining are toxic side effects on normal tissue. As this is unlikely to be fixed by fine-tuning the electrophilic warheads, the best option remains the application of predictive biomarkers, which can expand the therapeutic window for clinical success.

## Acknowledgement

The authors would like to thank the DKFZ Genomics and Proteomics Core Facility for support, expert advice and RNA sequencing and the DKFZ HUSAR server platform for providing the storage space and computing capacity to execute this project. We thank the Champions Oncology Team and Sebastian Brabetz (Champions Oncology, USA) for running the *ex vivo* PDX experiment, providing the PDX RNA sequencing data, and critical review of the manuscript. We thank the Helmholtz Validation Fund for supporting this work as part of the HOPE project.

## Materials and Methods (supplementary)

### CellTiter-Glo 3D cell viability assay and determination of EC50

Cells (H1793, A549, H661, H522) were seeded in a Nunclon™ Sphera™ 96-well u-shaped-bottom microplate (Thermo Scientific™, Cat No. 174925) at a density of 2,500 cells/100 µL/well in media containing 10 % FBS (Capricorn scientific), 100 U/mL penicillin, 100 µg/mL streptomycin (Sigma-Aldrich) and centrifuged for 10 min at 400 g. After 72 h, compact spheroids were treated with test compound (all concentrations in triplicate) for 24 h and viable cells were quantified in a FLUOstar OPTIMA ELISA reader 30 min after CellTiter-Glow 3D staining (Promega, Cat No. G9681). Mean values +/-SD were calculated. EC50 were calculated from dose response curves by GraphPad Prism.

### RNA isolation and RT-qPCR

Total RNA was isolated using NucleoSpin® RNA Kit according to the protocol (Macherey-Nagel, Cat No. 740955.250). Purified RNA was measured by Nanodrop (Thermo Fisher) and reverse transcription was performed using 1000 ng total RNA using Revert Aid First Strand cDNA Synthesis Kit (Thermo Scientific, Cat No. K1622) and PCR reactions with respective primer pairs were analysed in technical triplicates. Primer sequenes used for quantitative real-time PCR analyses are listed in Supplementary Table S5. GAPDH was used for normalization of gene expression.

### Cloning of sgRNA vectors for lentiviral particle production

Vectors were cloned, containing three individual small guide RNAs (sgRNA) for the genes *GSR*, *TXN*, *UGDH*, *GCLM*, *CBR1*, *PTRG1*, as well as two non-targeting sgRNAs. Sense and antisense oligo pairs of the sgRNAs (Supplementary Table S6) were phosphorylated and hybridized using a thermocycler (45 min 37 °C, 95 °C 4 min, cool down to 25 °C with a ramp rate of − 0.1 °C / sec): 1 µL sense oligo (100 µM, TE), 1 µL antisense oligo (100 μM, TE), 1 µL T4 DNA-ligase buffer (10 x), 0.5 µL T4 polynucleotide kinase, 6.5 µL H_2_O.

To obtain a vector backbone for sgRNA ligation, pXPR_502 (Addgene, Cat No. 96923) was digested using BsmBI (2 µg vector, 2 µL BsmBI-v2 (NEB, Cat No. R0739S), 2.5 µL 10x NEB3.1, up to 25 µL H_2_O). Mixture was incubated at 55 °C for 2 h 45 min and was then separated in a 1 % agarose gel (100 V, 45 min). Digestion results in a dropout of 30 bp, the remaining backbone (8664 bp) was extracted from the gel (Zymoclean gel DNA recovery kit, ZYMO RESEARCH, Cat No. D4007). A 1:200 dilution of the hybridized sgRNA oligos and the digested backbone were ligated at RT for 1 h: 50 ng digested and purified pXPR_502 plasmid, 1 µL phosphorylated and annealed oligo pair (1:200 dilution in H_2_O), 2 µL T4 DNA-Ligase Buffer (10 x), 0.25 µL T4 DNA-Ligase, up to 20 µL H_2_O.

The ligation mix was then used to transform NEB® Stable Competent E. coli (NEB, Cat No. C3040H). Therefore, bacteria were thawed on ice for 10 min, the ligation mix was added, and the cells were incubated for 30-60 min on ice. Next, a heat shock at 42 °C for 45 sec was performed, followed by incubation on ice for 2 min. Then, 250 µL SOC medium was added and the cells were incubated at 37 °C for 1 h to recover. The entire vial of bacteria was plated in ampicillin agar plates (100 µg/mL) and incubated for 12-16 h at 37 °C. The next day, single colonies were picked and used to inoculate small liquid cultures (3 mL LB-Medium, 100 µg/mL ampicillin), incubation at 37 °C for 12-16 h, 220 rpm. Using a miniprep kit (NucleoSpin Plasmid, Macherey-Nagel, Cat No. 740588.250), the vectors were isolated.

To validate the correct insertion, plasmids were sequenced (LightRun Tubes, eurofins) using a U6 sequencing primer (5’-GAGGGCCTATTTCCCATGATTCC-3’). Correct clones were then used for bigger liquid cultures (150 mL LB-Medium, 100 µg/mL ampicillin) and vectors were isolated using a HiSpeed Plasmid Midi Kit (Qiagen, Cat No. 12643).

### Lentivirus particle production and transduction of target cells

Polyclonal HCC827and H23 cells constitutively expressing the CRISPR activation machinery were engineered by transducing wild-type cells with lentiviral particles carrying a dCas9-VP64 (lenti dCas9VP64_Blast, Addgene plasmid #61425) at a multiplicity of infection (MOI) of ∼0.5. dCas9VP64 expressing cells were then transduce with pXPR_502 carry the sgRNA and transcriptional activation domains p65-HSF1. CRISPRa cell lines were selected for puromycin (4 µg/µL for sgRNAs) and blasticidin (25 µg/µL for dCas9-VP64 construct) every 2-3 weeks for 72 h. Large-scale lentivirus production was performed using a second-generation lentivirus system and a calcium phosphate transfection kit (Invitrogen, Cat No. K278001) in HEK293T cells. Briefly, early passaged HEK293T cells were co-transfected with the lentiviral transfer plasmid, a packaging plasmid (psPAX2, Addgene plasmid #12260), as well as with a plasmid encoding the VSV-G envelope (pMD2.G, Addgene plasmid #12259). Viral supernatant was collected 26-30 h post transfection and stored at −80 °C until use. All experimental procedures for lentivirus production and transduction were performed in a biosafety level 2 laboratory.

### Lipid peroxidation

As an indicator of ferroptosis, lipid peroxidation was analyzed in cells stained with Bodipy 581/591 C11 (Invitrogen). Cells were stained with 1.5 µM Bodipy 581/591 C11 diluted in the culture medium for 2 h at 37 °C. Afterwards, cells were detached with TrypLE (Gibco), washed with PBS, and analyzed using the flow cytometer BD LSR Fortessa (BD Biosciences). The ratio of the oxidized (excitation 488 nm, emission 530/30) and reduced (excitation 561 nm, emission 610/20) dye was calculated for each cell in the FlowJo software.

### Iron pool assay

FIP-1 probe (FRET Iron Probe) was a gift from Christopher Chang. It enables ratiometric fluorescence imaging of labile iron pools in living cells [73]. Cells were seeded on a 12-well plate to achieve 60-70 % confluence on the day of the assay. 250 μM deferoxamine (DFO) (Sigma) or PBS was added to Fluorobrite media containing 10 % FBS in wells containing cells and incubated at 37 °C for 6 h. After the incubation, media was aspirated and cells were washed with 500 μL HBSS. Then 500 μL HBSS containing 10 μM FIP-1 (diluted from 5 mM stock) was added to each well and this was incubated at 37 °C for 90 min. Cells were harvested by trypsinization, washed once with PBS and FIP-1 fluorescence was analyzed with a flow cytometer (Guava easyCyte 14HT, Luminex) in two channels: Green-B (“Green”, excitation 488, emission 512/18), which is high in the presence of iron, and Yellow-B (“FRET”, excitation 488, emission 575/25), which is low in the presence of iron. Mean Green/FRET ratio was obtained for each cell line, and the signal was normalized to the cells treated with an iron chelator - deferoxamine (DFO, final concentration 300 µM, 6 h treatment prior to the staining). Unstained cells were used as a control for flow cytometry.

### PDX model

*Ex vivo* PDX drug sensitivity assays were performed by Champions Oncology. Cryopreserved tumor fragments, derived from the NSCLC models CTG-2842 and CTG-0852 were thawed, mechanically dissociated and filtered through 400 µm and 200 µm pores. After 24 h recovery and viability assessment by CellTiter-Glo in low attachment plates, cells were plated into 384-well format and drug added. After 6 days, drug effects were assessed via CellTiter-Glo.

**Supplementary Figure S1.**
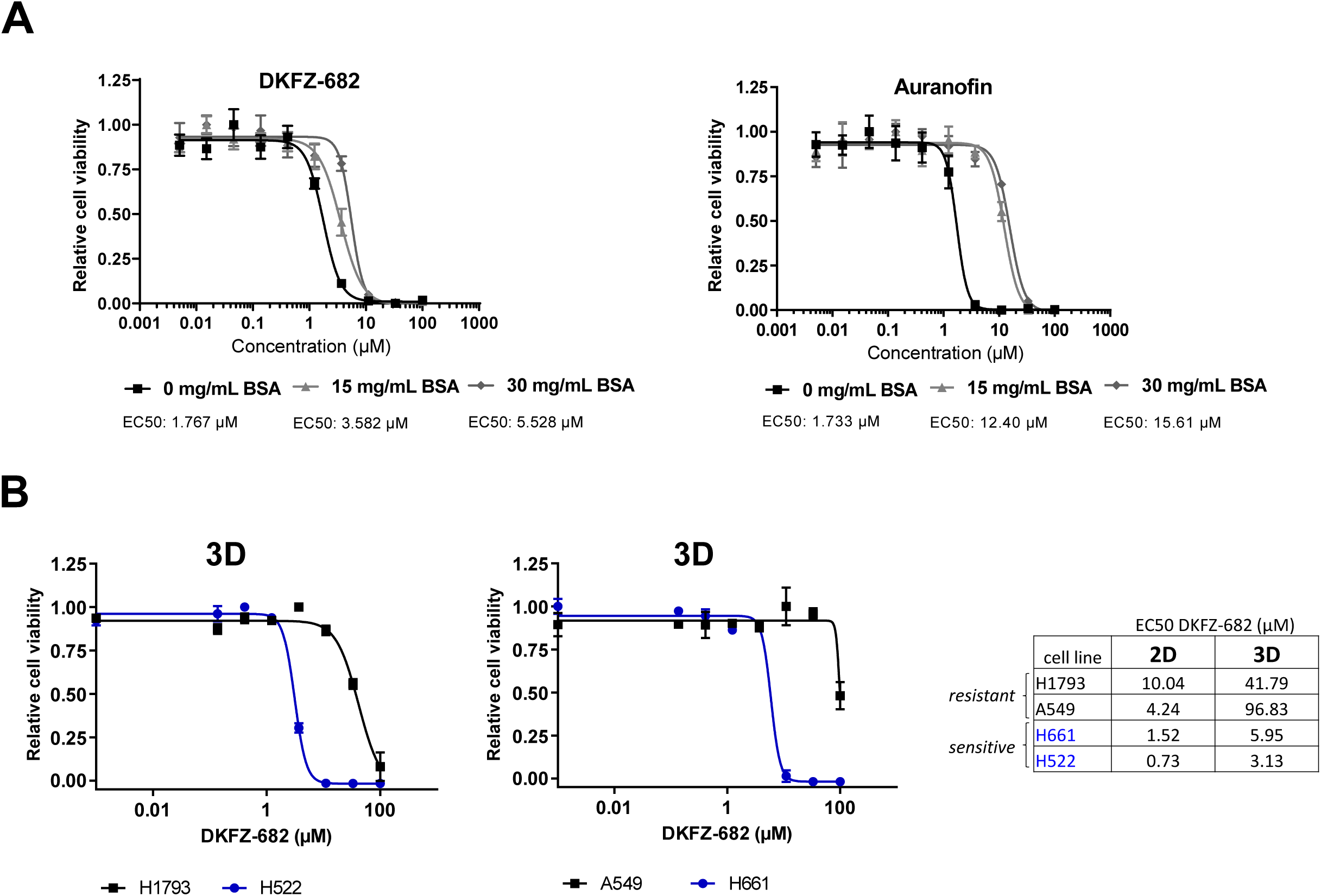
Inhibitory activities of DKFZ-682 and its analog Auranofin in cell culture assays. **(A)** Various concentrations of DKFZ-682 or auranofin were prepared in medium containing 0, 15 or 30 mg/mL additional BSA. After 1 h preparation of solutions, H838 cells were treated for 8 h in 96-well plates with the dilution series of each compound. After washing with fresh medium, inhibitor free medium was added. The numbers of surviving cells were quantified 64 h later using the CellTiter-Blue assay. EC50 values were determined from dose-response curves using GraphPad Prism. **(B)** Three-dimensional (3D) cell spheroids (H1793 and A549 resistant to DKFZ-682 shown in black; H661 and H522 sensitive to DKFZ-682 shown in blue) were treated with a concentration series of DKFZ-682 for 24 h and the cell viability was quantified by the CellTiter-Glow 3D assay. EC50 values were determined from dose-response curves using GraphPad Prism. The graph is representative of two independent experiments each performed in triplicates (error bars indicate SD). For comparison of DKFZ-682 activity in 2D monolayer and 3D spheroid model (right panel), 2D EC50 data of Fig. 1D were used.

**Supplementary Figure S2.**
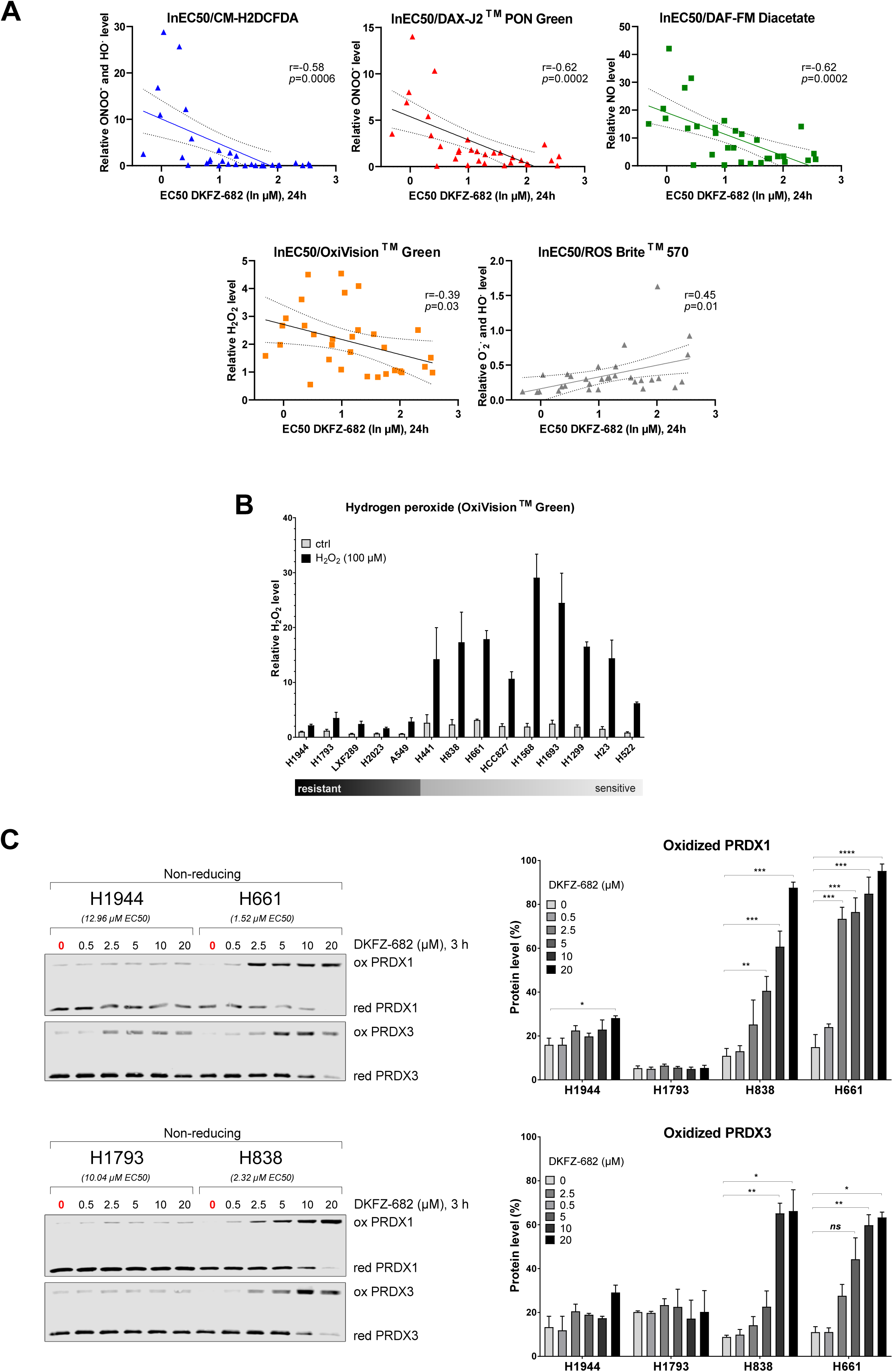
Redox buffer capacity of ROS drug resistant versus sensitive cells. **(A)** Scatter plot of DKFZ-682 EC50 (ln, µM) versus relative levels of basal ROS/RNS. Correlation is assessed by Pearson coefficient on n=31 NSCLC cell lines. NSCLC cell lines were treated with a concentration series of DKFZ-682 for 24 h and the cell viability was quantified by the CellTiter-Blue assay. EC50 values were determined from dose-response curves using GraphPad Prism. For ROS/RNS detection cells were stained with CM-H2DCFDA (ONOO^-^, HO^•^), DAF-FM Diacetate (NO), DAX-J2™ PON Green (ONOO^-^), OxiVision^TM^ Green peroxide sensor (H_2_O_2_) or ROS Brite^TM^ 570 (O_2_^•-^, HO^•^) fluorescent dyes and analysed by flow cytometry. **(B)** NSCLC cells were stained with OxiVision^TM^ Green peroxide sensor for 20 min and treated with or without H_2_O_2_ (100 µM) for further 10 min without medium change, and analysed by flow cytometry. The graphs summarize the relative data of independent experiments (n=3-4, error bars indicate SEM). **(C)** Cells were incubated with the indicated concentration of DKFZ-682 for 3 h. Oxidized (ox) and reduced (red) status of PRDX1 and PRDX3 proteins were analysed by immunoblotting. Bar diagrams summarize the quantitative results from independent experiments (n=2-3, error bars indicate SD; \**q*<0.05, \*\**q*<0.01, \*\*\**q*<0.001, \*\*\*\**q*<0.0001, two-tailed unpaired *t* test using the original data).

**Supplementary Figure S3.**
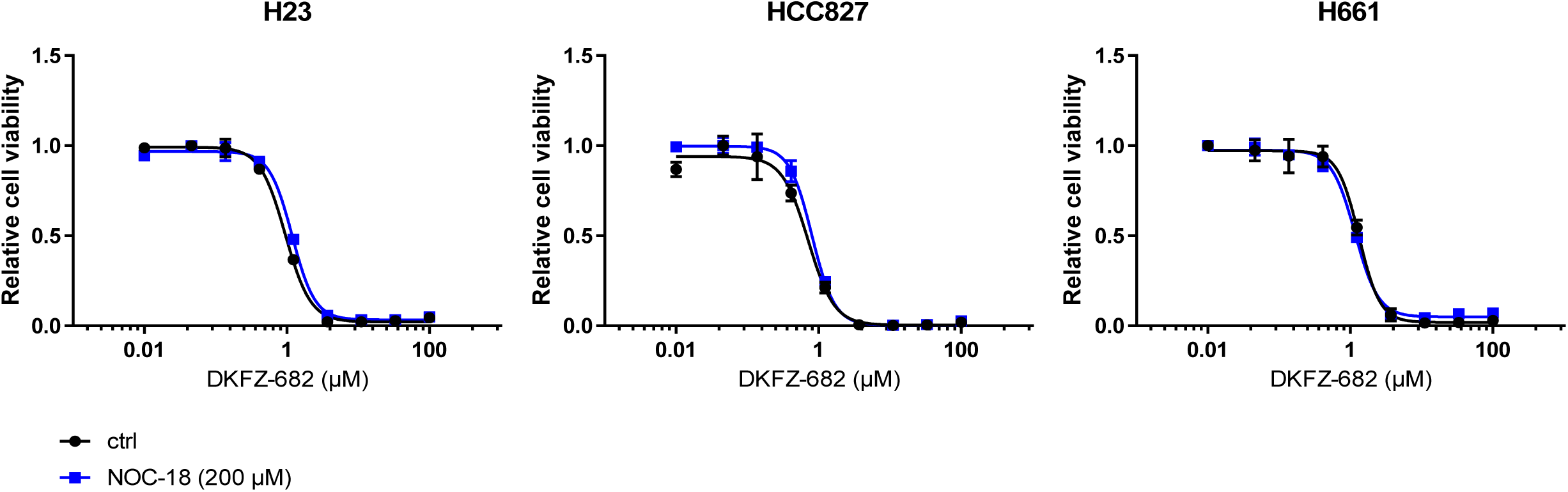
Nitric oxide (NO) increase does not reduce DKFZ-682 toxicity. H23, H661 and HCC827 cells were treated with a concentration series of DKFZ-682 for 24 h in the presence of NO donor NOC-18 (200 µM), and the cell viability was quantified by the CellTiter-Blue assay. Data points represent results from three technical replicates. The results are representative of two independent experiments.

**Supplementary Figure S4.**
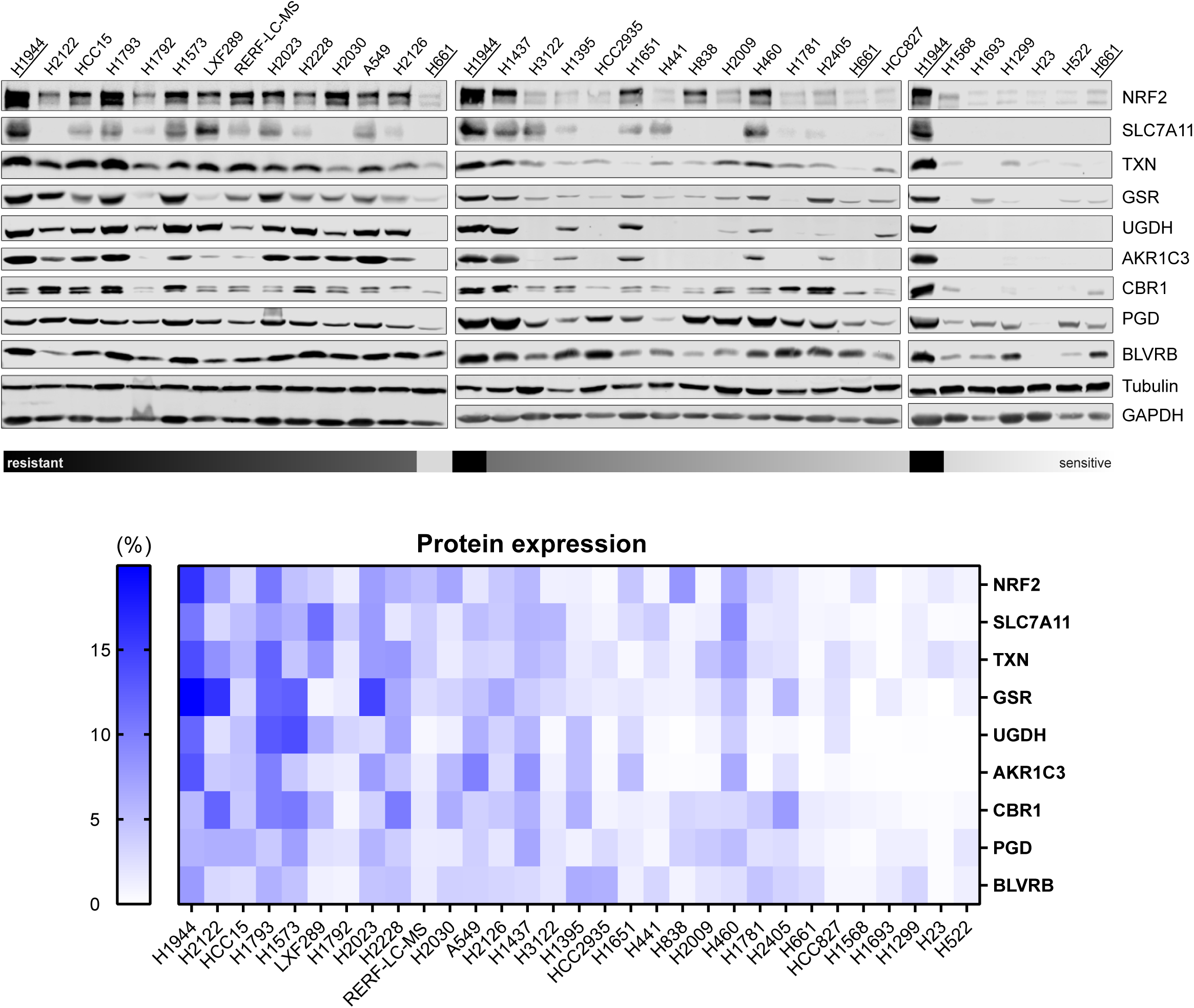
ACB protein expression in NSCLC cell lines. Protein levels of NRF2 and different ACBs in 31 NSCLC cell lines were analysed by immunoblotting. Heatmap with the expression values of NRF2 and 8 ACB genes. First protein expression in H1944 was set to 1 for each gene and for each experiment (n=2-6). Then the data were normalized for each gene separately. The smallest value of row was set to 0 %, and the sum of all values in the row was set to 100 %.

**Supplementary Figure S5.**
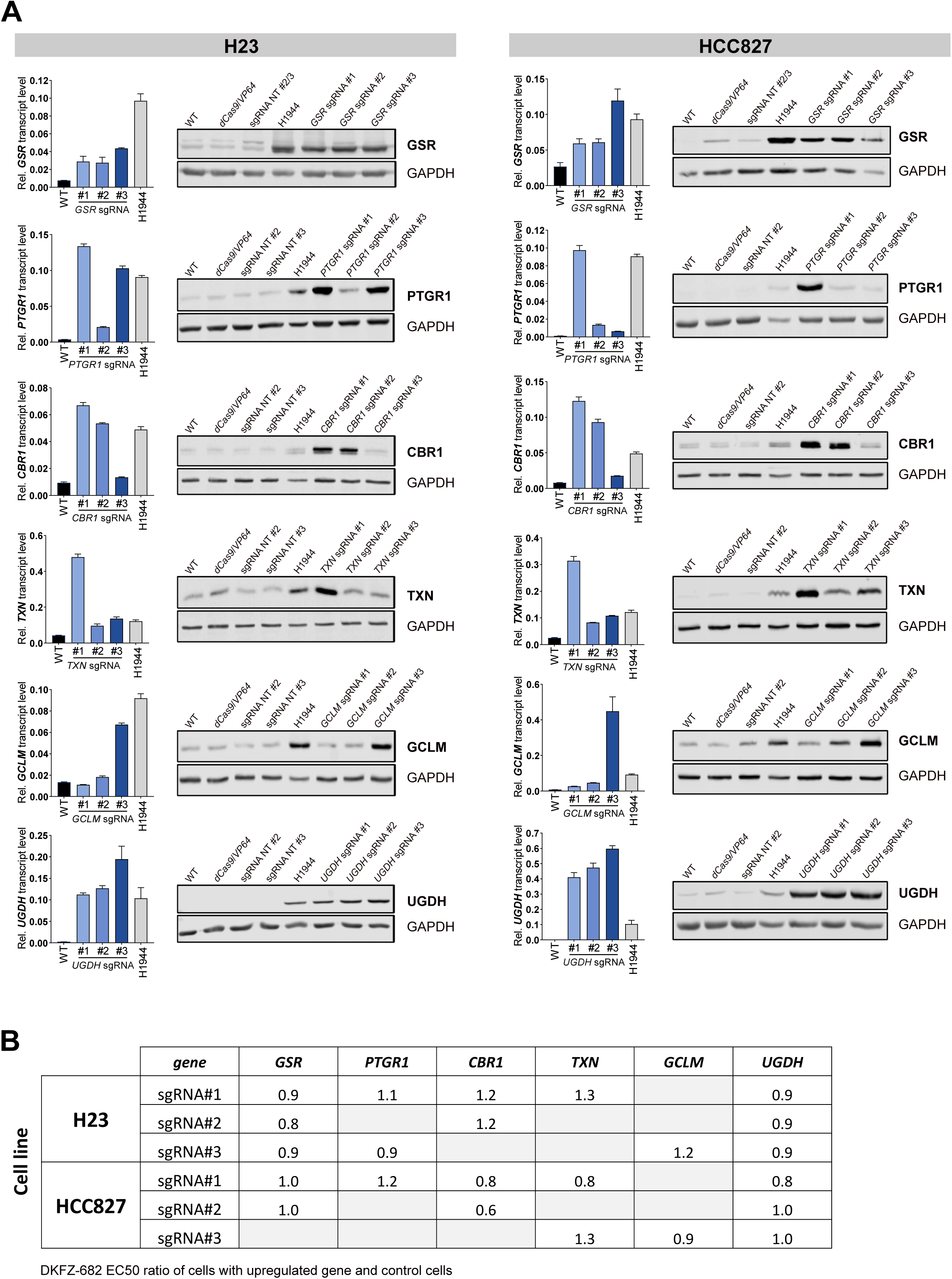
Overexpression of single ACB genes using CRISPR activation technology. **(A)** ACB transcript and protein levels in H23 and HCC827 wild type (WT) or CRISPRa cell lines expressing an enzymatically inactive Cas9 protein (dCas9, only RNA binding activity) with linked transcriptional activators VP64 alone (dCas9/VP64), or with target gene specific (sgRNA) or non-targeting (NT sgRNA) small guide RNAs were analysed by qPCR and immunoblotting respectively. Bars represent mean ± SD of three technical replicates. Transcript level of each gene was normalized to *GAPDH* level. Protein and transcript levels of H1944 were used as control. **(B)** H23 and HCC827 CRISPRa cell lines with non-targeting (NT sgRNA) small guide RNAs or overexpressing *GSR*, *PTGR1*, *CBR1*, *TXN*, *GCLM* or *UGDH* (target gene sgRNA) were treated with a concentration series of DKFZ-682 for 24 h and the cell viability was quantified by the CellTiter-Blue assay. Three technical replicates were performed. EC50 values were determined from dose-response curves using GraphPad Prism. The results of DKFZ-682 EC50 ratio of cells with upregulated gene (target gene sgRNA) and control cells (NT sgRNA) are shown in the table.

**Supplementary Figure S6.**
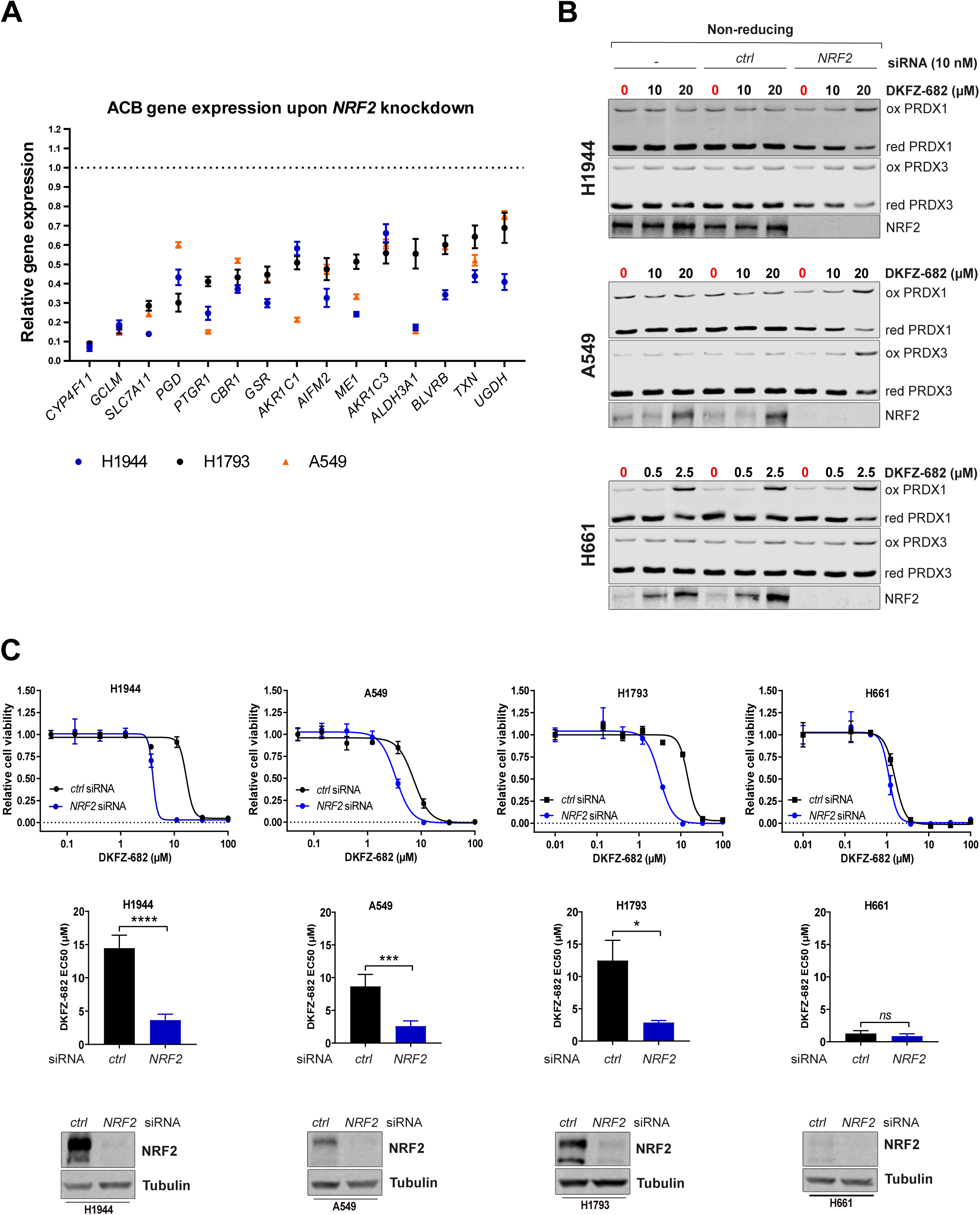
Knockdown of *NRF2* decreases ROS protection efficiency of DKFZ-682 resistant cells. **(A)** H1944 and H1793 cell lines were transfected with nonsense or *NRF2* specific siRNA. The ACB gene expression 48 h after siRNA transfection was quantified using the Affy Clariom S Human array. Expression data derived from *NRF2* knockdown in A549 cells are derived from GSE38332 (GEO accession number). Fold change reduction was calculated as a ratio of three biological replicates each of cells transfected with nonsense or *NRF2* siRNA. The dotted line indicates relative ACB expression in control cells transfected with nonsense siRNA. **(B)** Cell lines were transfected with nonsense (*ctrl*) or *NRF2* siRNA for 48 h and then treated with the indicated concentration of DKFZ-682 for 3 h. Oxidized (ox) and reduced (red) levels of PRDX1 and PRDX3 proteins were analysed by immunoblotting. Representative western blots of at least 2 independent experiments are shown. **(C)** H1944, A549, H1793 and H661 cell lines were transfected with nonsense (ctrl) or *NRF2* specific siRNA. After 48 h cells were treated with a concentration series of DKFZ-682 for 24 h and the cell viability was quantified by the CellTiter-Blue assay. Bar diagrams show the mean of EC50 data from independent experiments (n=4 H1944 and A549, n=2 H1793 and H661) each performed in triplicates (error bars indicate SD, \**q*<0.05, \*\*\**q*<0.001, \*\*\*\**q*<0.0001, two-tailed unpaired *t* test). NRF2 protein expression was analysed by immunoblotting. Representative western blots are shown.

**Supplementary Figure S7.**
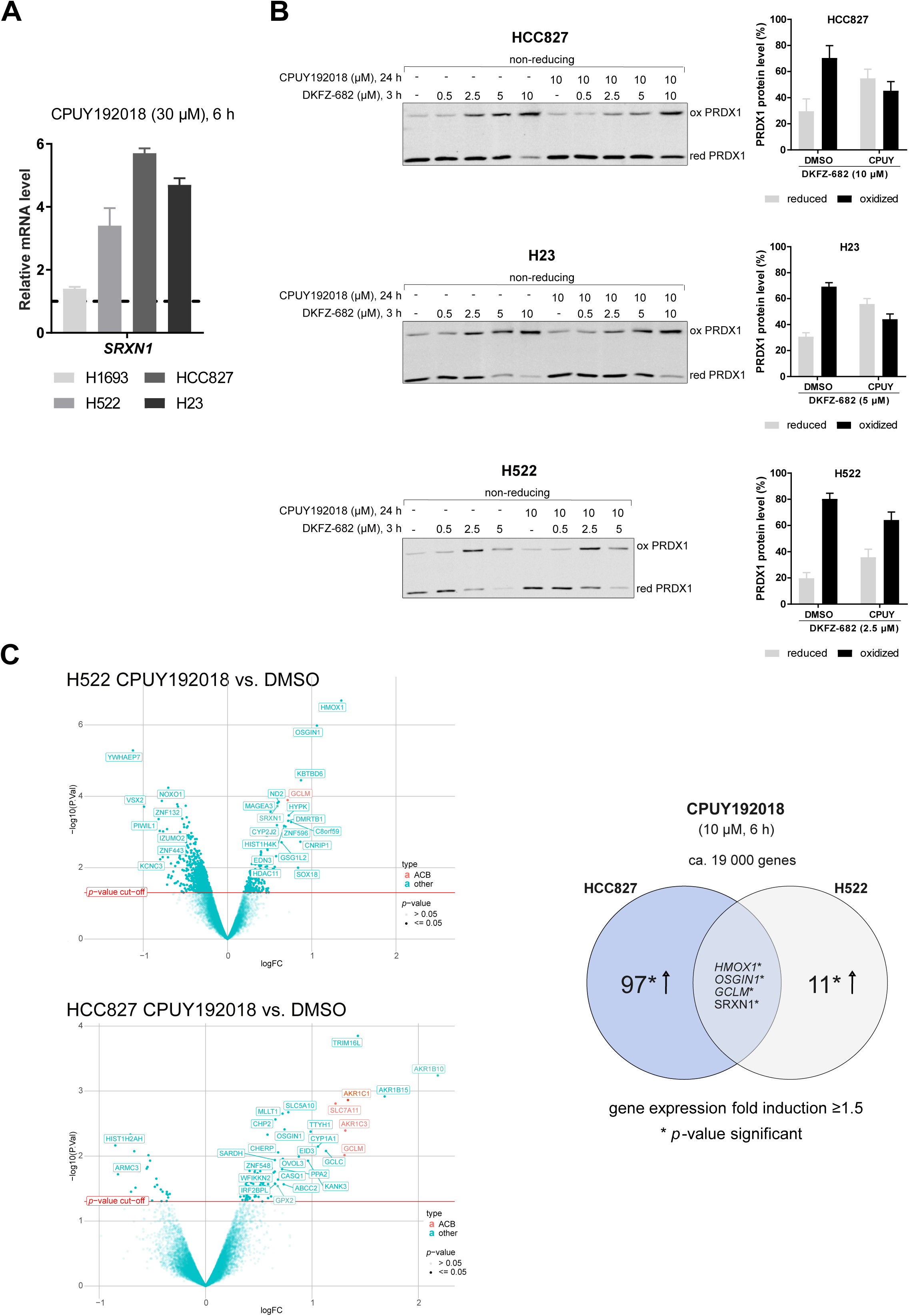
An impact of NRF2 overexpression in DKFZ-682 sensitive cells on redox buffer capacity. **(A)** HCC827, H522, H23 and H1693 cell lines were treated with CPUY192018 (30 µM) or DMSO for 6 h. *SRXN1* transcript level analysis was performed using qPCR assay. *SRXN1* expression in DMSO treated cells was set to 1. Relative data represent mean of independent experiments each performed in triplicates (n=2, error bars indicate SD). **(B)** HCC827, H23 and H522 cell lines were treated with DMSO or CPUY192018 (10 µM). After 24 the cell medium was changed and cells were treated with DKFZ-682 for 3 h. Oxidized (ox) and reduced (red) levels of PRDX1 protein were analysed by immunoblotting. Bar diagrams show the quantitative results only for prominent DKFZ-682 concentration different for each cell line (n=2, error bars indicate SD). **(C)** Volcano plot (left panel) of genes significantly up-or downregulated in H522 and HCC827 cell lines after 6 h treatment with CPUY192018 (10 µM) or DMSO. Transcript data analysis using RNAseq. An overview of differentially expressed ACB genes under CPUY192018 application in HCC827 and H522 cell lines (right panel). Results are representative of three independent experiments each performed in triplicates (error bars indicate SD, \**q*<0.05, unpaired multiple *t* test, comparison DMSO versus drug treatment).

**Supplementary Figure S8.**
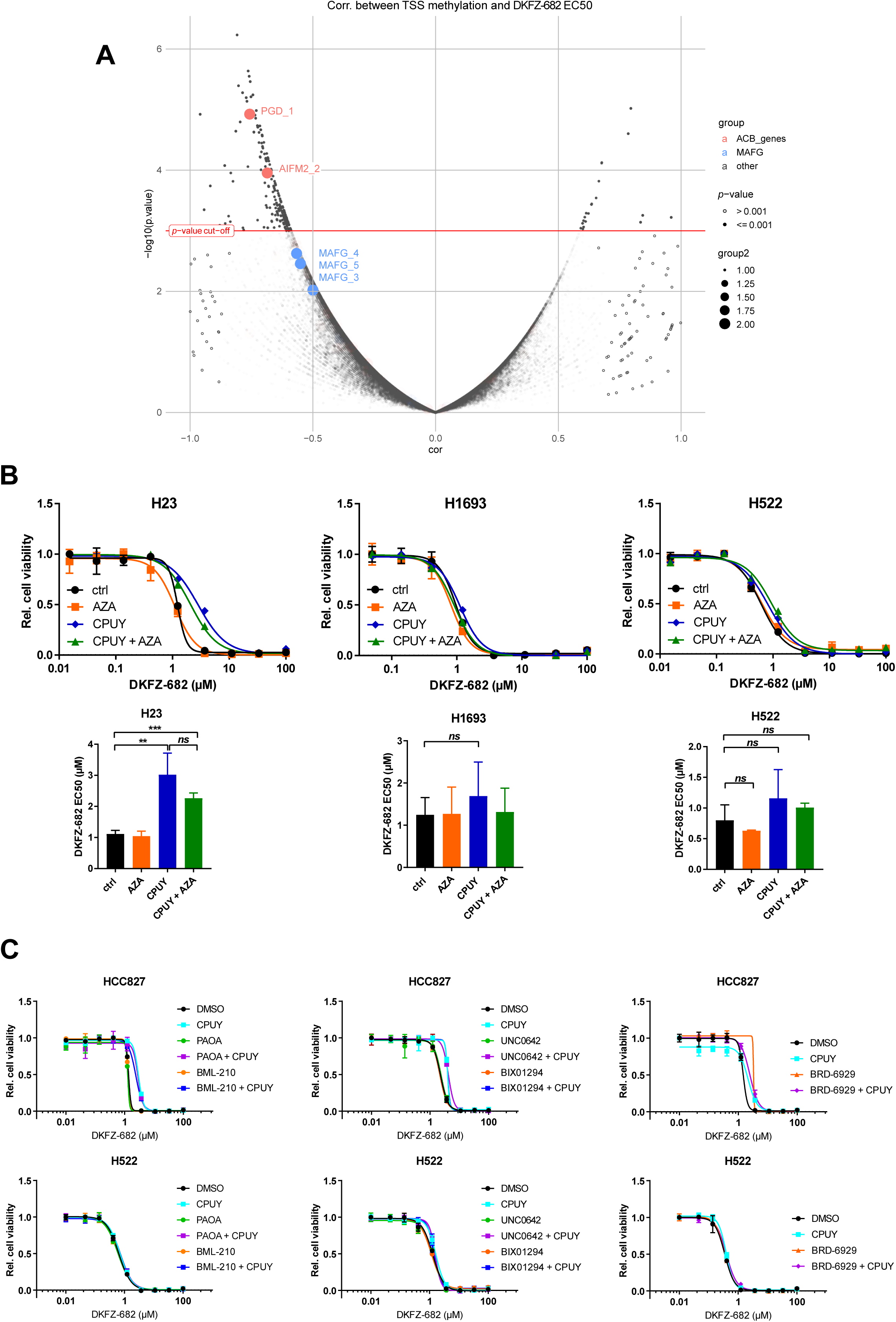
ACB expression remains repressed after treatment with DNMT-, HDAC- or HMT inhibitors. **(A)** Volcano plot shows the results of correlation analysis between methylation values of transcription start sites CpG clusters (reduced-representation bisulfite sequencing data) and EC50. **(B)** H23, H1693 and H522 cells were treated with CPUY192018 (CPUY, 10 µM) after 3 days of culture with or without azacytidine (AZA, 2.5 µM) added on day 1 and 2 after seeding (day 0). After 24 h, cells were treated with a range of concentrations of DKFZ-682 for 24 h. Cell viability was assessed using CellTiter-Blue assay. Bar diagrams represent mean ± SD of independent experiments (n=2-3, \*\**q*<0.01, \*\*\**q*<0.001, *ns*, not significant, paired Student *t* test using the original data). **(C)** HCC827 and H522 cells were treated for 24 h with HDAC inhibitors (5 µM BML-210, 5 µM PAOA or 5 µM BRD-6929) or for seven days with G9a inhibitors (2 µM UNC0642 or 2 µM BIX01294). After this period, CPUY192018 (10 µM) was added for 24 h. Next, cells were treated with series dilutions of DKFZ-682, and after 24 h, cell viability was quantified by the CellTiter-Blue assay. Bars represent results from three technical replicates.

**Supplementary Figure S9.**
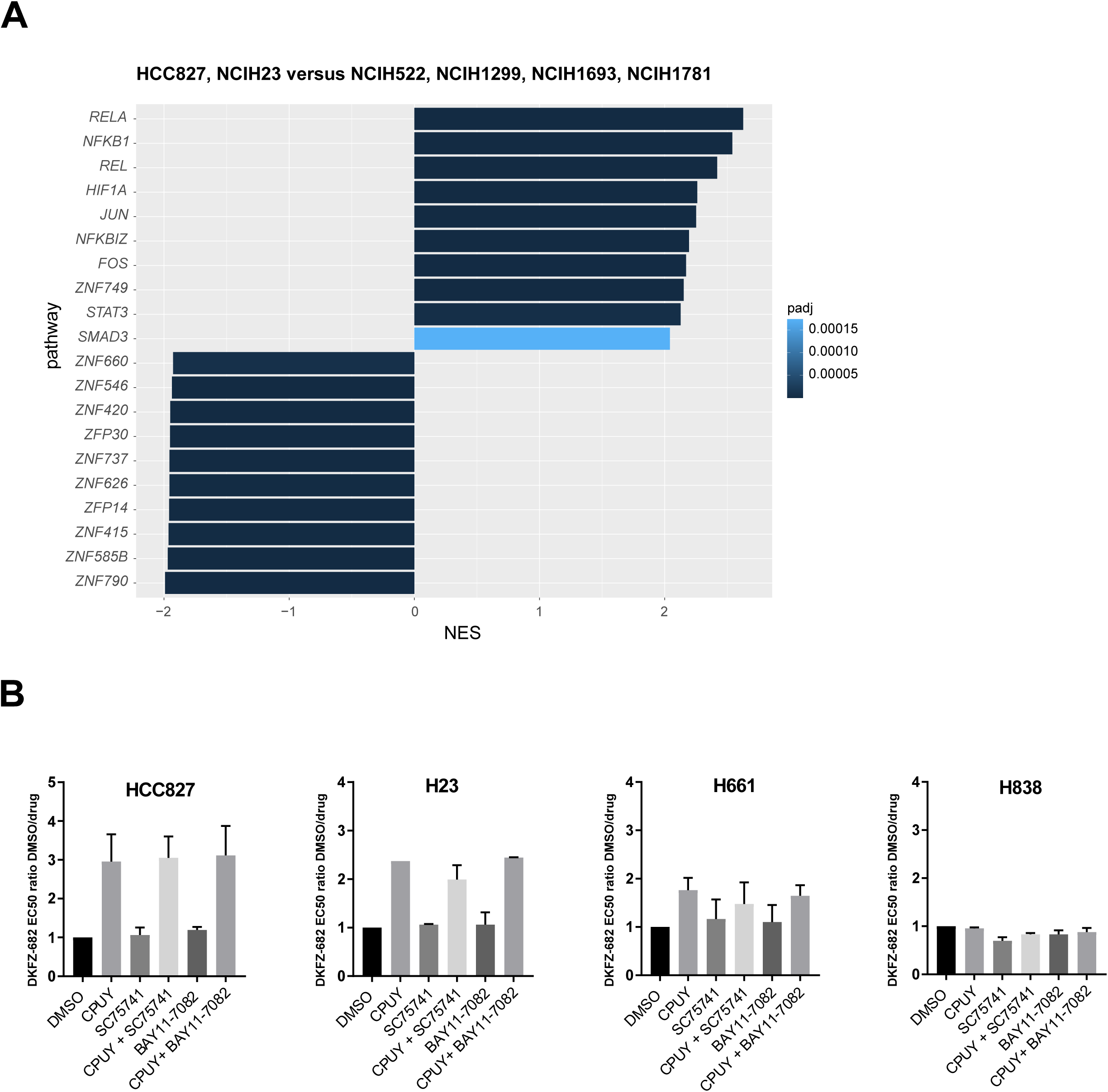
NFKB is not required for NRF2-mediated induction of drug resistance. **(A)** Gene set enrichment analysis. We performed differential expression analysis between two groups - HCC827, H23 and H522, H1299, H1693, H1781 using limma Bioconductor package. Then, with differential expression results we performed GSEA analysis using as genes sets “transcription factor – genes” association data from Dorothea Bioconductor package. We plotted 10 pathways with highest positive NES and 10 pathways with lowest negative NES from GSEA results. **(B)** Cells were pretreated with DMSO as a control, CPUY192018 (CPUY, 10 µM), SC75741 (1 µM), BAY11-7082 (1 µM), or in combination of CPUY192018 with SC75741 or with BAY11-7082. After 24 h cells were treated with a concentration series of DKFZ-682 for 24 h and the cell viability was quantified by the CellTiter-Blue assay. The results of DKFZ-682 EC50 ratio of DMSO and drug treated cells are shown. Bar diagrams summarize the quantitative results of independent experiments (n=2, error bars indicate SD) each performed in triplicates.

**Supplementary Figure S10.**
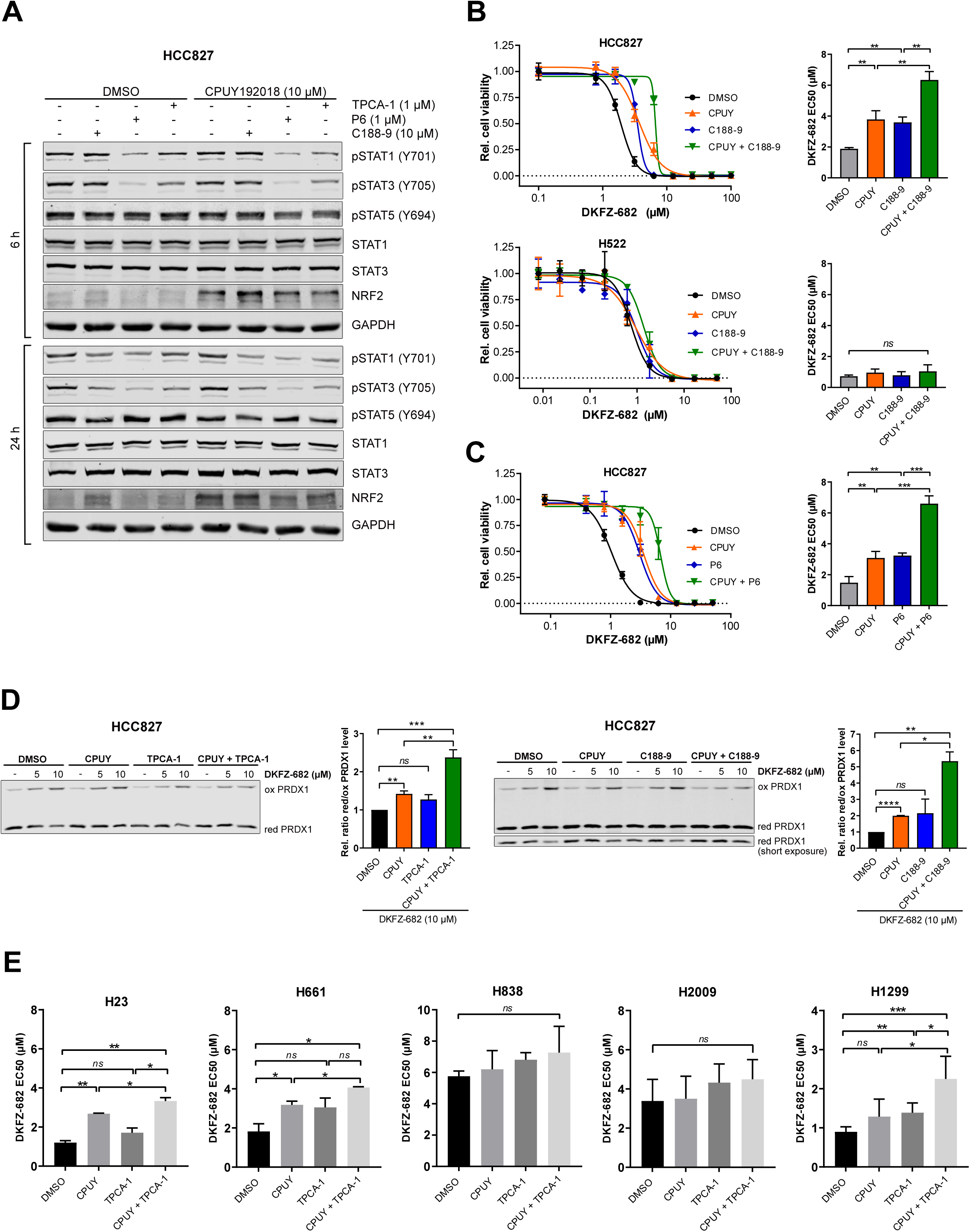
Inhibition of STAT activity enhances NRF2 dependent drug resistance. **(A)** HCC827 cells were treated with DMSO as a control, TPCA-1 (1 µM), P6 (1 µM) und C188-9 (10 µM) alone or in the presence of CPUY192018 (10 µM) for 6 and 24 h. STAT protein expression was analysed by immunoblotting. Representative western blots of two independent experiments are shown. **(B-E)** Cells were treated with DMSO as a control, 10 µM C188-9 **(B, D)**, 1 µM P6 **(C),** 1 µM TPCA-1 **(D, E)**, alone or in the presence of CPUY192018 (CPUY, 10 µM) for 24 h. **(B, C)** After pretreatment with indicated compounds cells were treated with series dilutions of DKFZ-682, and after 24 h, cell viability was quantified by the CellTiter-Blue assay. **(D)** Then the cell medium was changed and cells were treated with DKFZ-682 (5 µM or 10 µM) for 3 h. Oxidized (ox) and reduced (red) levels of PRDX1 protein were analysed by immunoblotting. Ratio of oxidized and reduced PRDX1 in control cells was set to 1. Bar diagrams show the mean of EC50 data of independent experiments (n=3 **(B-D)**, n=2-3 **(E)**) each performed in triplicates (error bars indicate SD (**B, C, E**) or SEM (**D**), \**q*<0.05, \*\**q*<0.01, \*\*\**q*<0.001, *ns*, not significant, two-tailed unpaired *t* test).

**Supplementary Figure S11.**
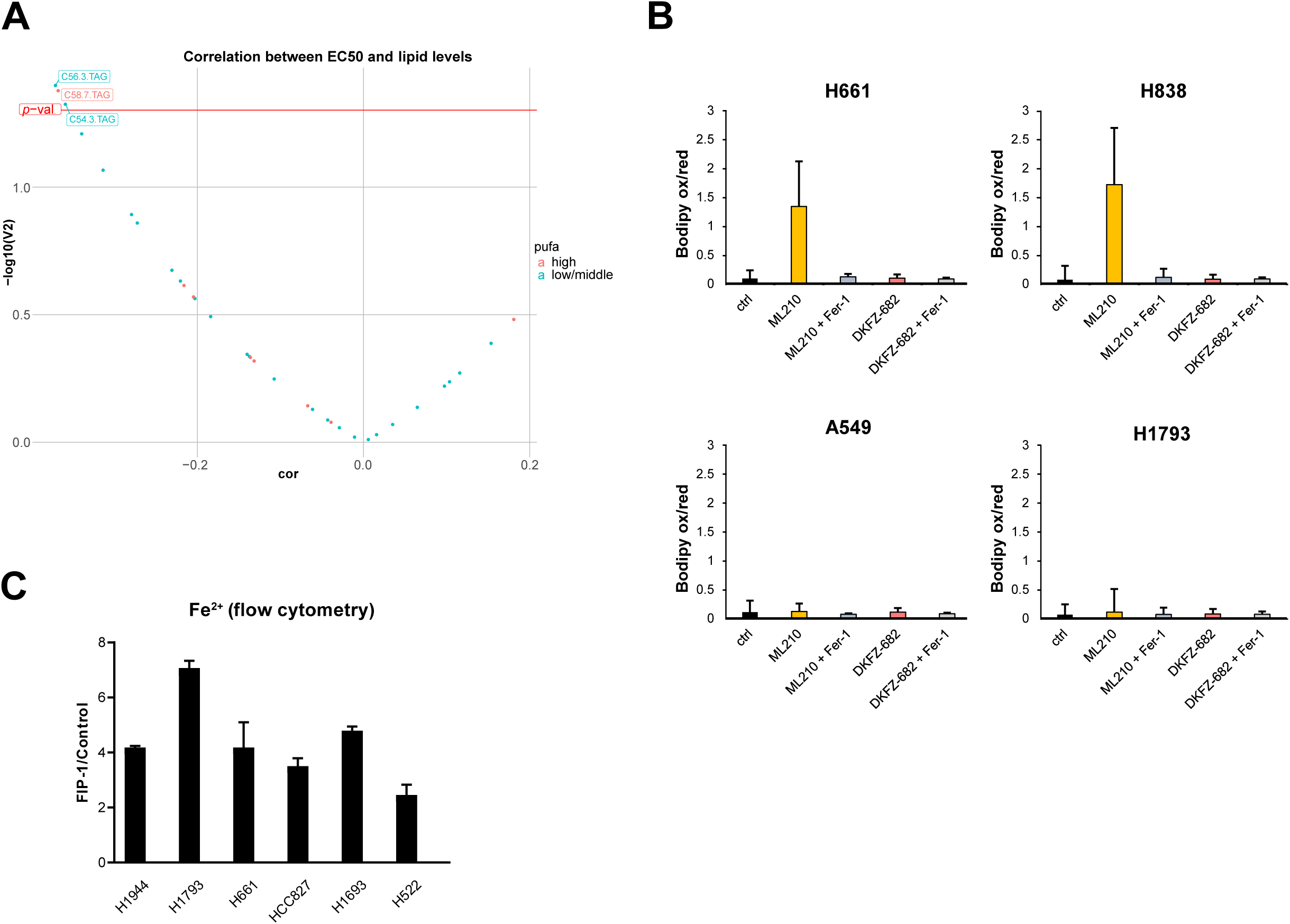
Lipid composition and steady state iron content does not correlate with sensitivity to DKFZ-682. **(A)** Volcano plot visualizing correlations between lipid levels and EC50 values. Liquid chromatography–mass spectrometry metabolite data. **(B)** DKFZ-682 does not induce ferroptosis, as shown by the lipid peroxidation sensor Bodipy 581/591 C11. After a pre-treatment with ferroptosis inhibitor ferrostatin-1 (Fer-1, 10 µM), cells were stained with Bodipy (3 µM, 3 h) in the presence of either DKFZ-682 (5 µM for H661 and H838, 20 µM for A549 and H1793) or a ferroptosis inducer ML210 (10 µM). After washing, cells were analyzed using a flow cytometer. The ratio of the oxidized (excitation 488 nm, emission 530/30) and reduced (excitation 561 nm, emission 610/20) dye is shown for each condition. **(C)** Flow cytometry labile iron pool assay in the NSCLC panel with FRET Iron Probe 1 (FIP-1) enables ratiometric fluorescence imaging of labile iron pools in living cells. Mean Green/FRET ratio was obtained for each cell line, and the signal was normalized to the cells treated with an iron chelator - deferoxamine (DFO). No significant difference was observed between high and low ACB expressing cell lines. Error bars denote SD, n=3.

**Supplementary Figure S12.**
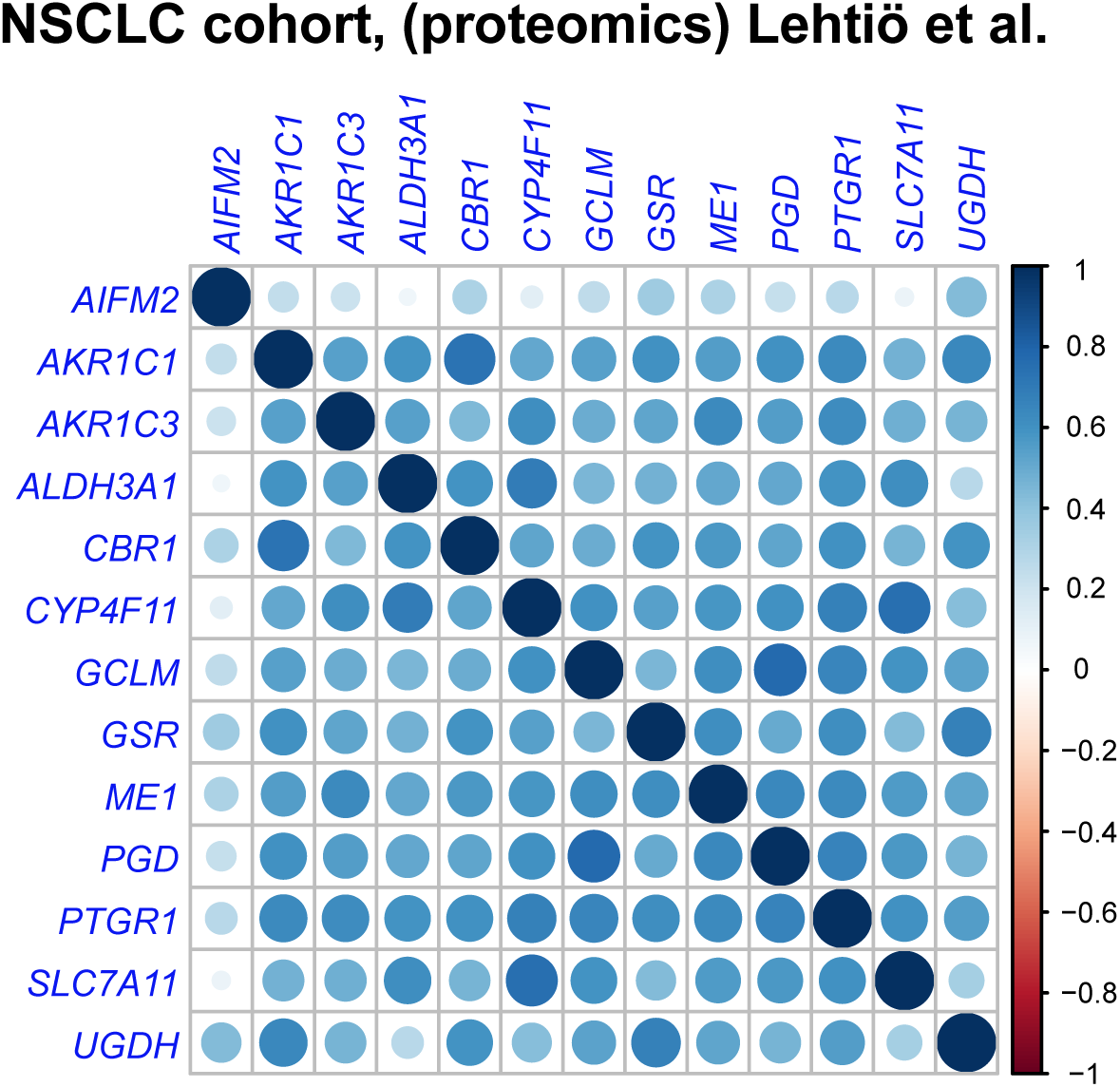
Coordination expression of ACB proteins in NSCLC tumors. Correlation plots show protein to protein correlations for ACB genes in 141 NSCLC tumors, derived from proteomics data Lehtio et al. [35].

**Supplementary Figure S13.**
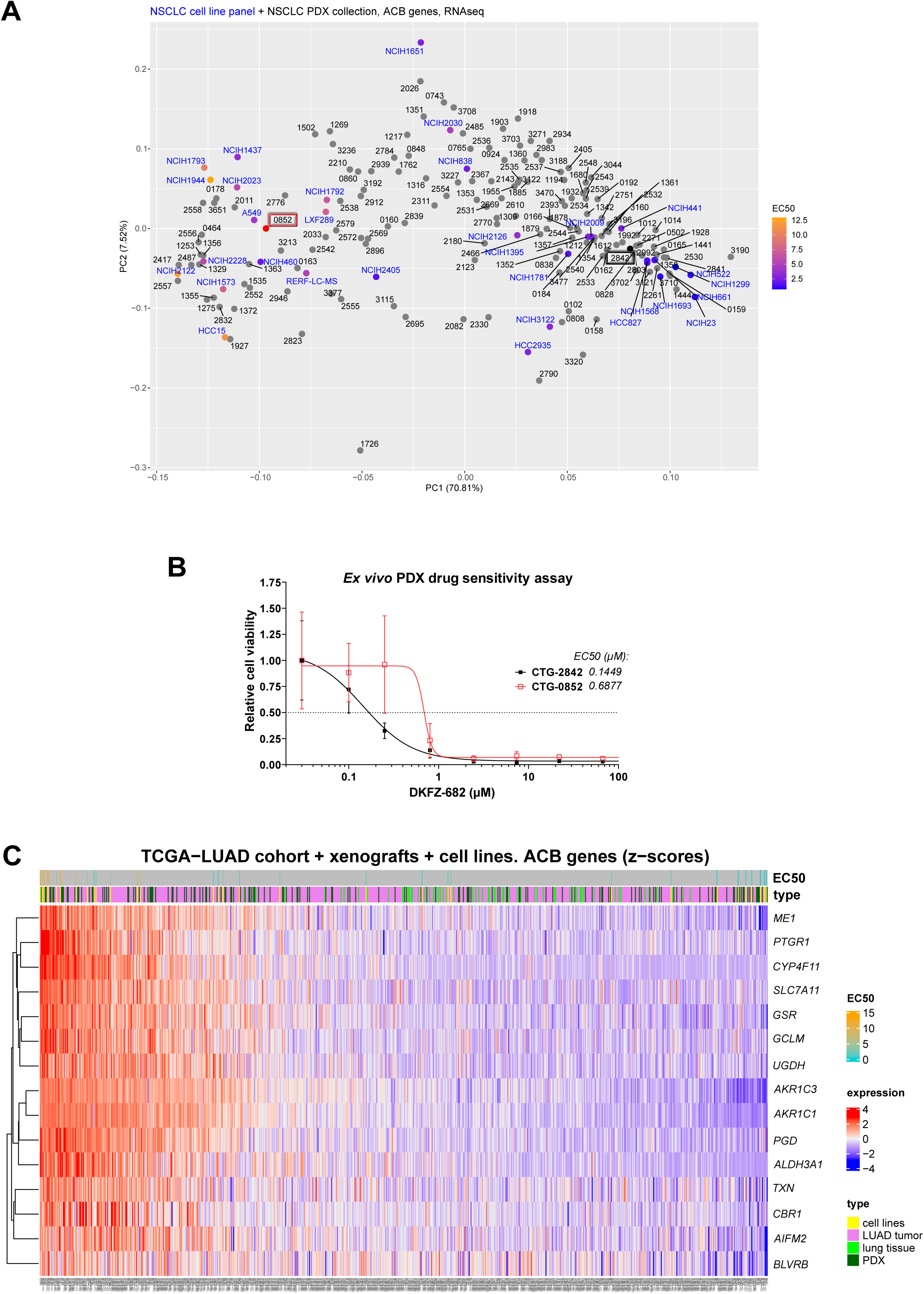
Expression space of ACBs in cell lines, PDX models and LUAD patients. **(A)** Principal component analysis plot (1^st^ and 2^nd^ PCA components) on ACB genes. Samples include NSCLC cell lines (blue) and PDX (black) models. Sensitivity of cell lines is indicated by the color code (blue – sensitive, orange – resistant). **(B)** Cell killing effect after 5 days incubation of tumor fragments (black – low ACBs, red – high ACBs) with DKFZ-682. Cell viability was quantified with CellTiter-Glo. Data are presented as the mean ± SD of six technical replicates. **(C)** Heatmap with ACB genes expression in cell lines TCGA-LUAD tumor and control samples, and PDX models. Rows (genes) are clustered using “complete linkage” method and “Euclidean” distance. Top annotation includes EC50 values (for cell lines) and sample type.

**Supplementary Figure S14.**
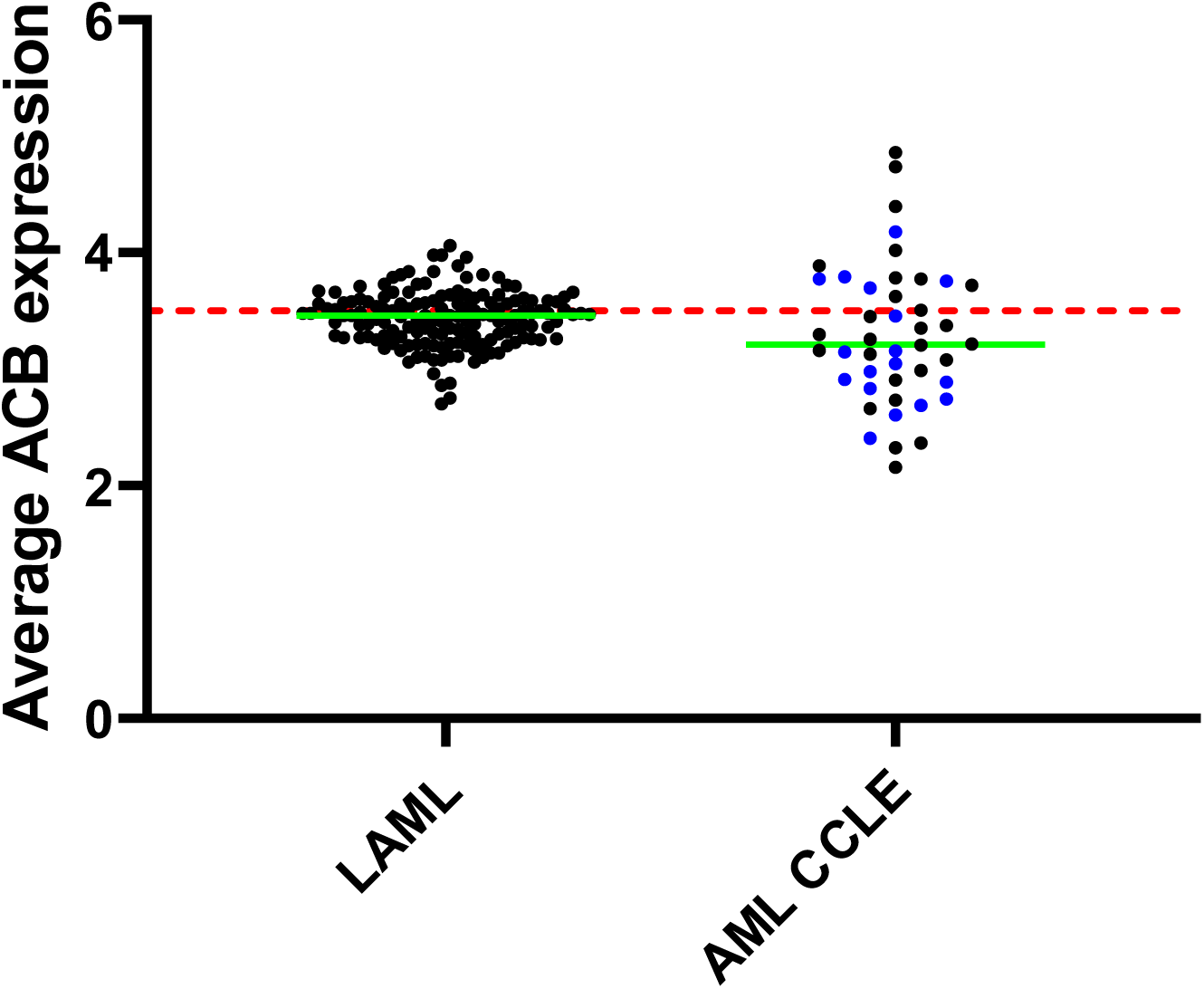
Comparison of average ACB expression in AML samples from the TCGA dataset and AML cell lines from CCLE. The red dashed line indicates ACB expression in the NSCLC line H23. Blue symbols show cell lines included in the AML cell panel used to determine the activity profile of DKFZ-682. Green lines represent the mean value for each dataset. Cell line data are in log_2_(TPM+1) units. TCGA-LUAD expression data were converted from RPKM to log_2_(TPM+1) units.

**Supplementary Table S1:**
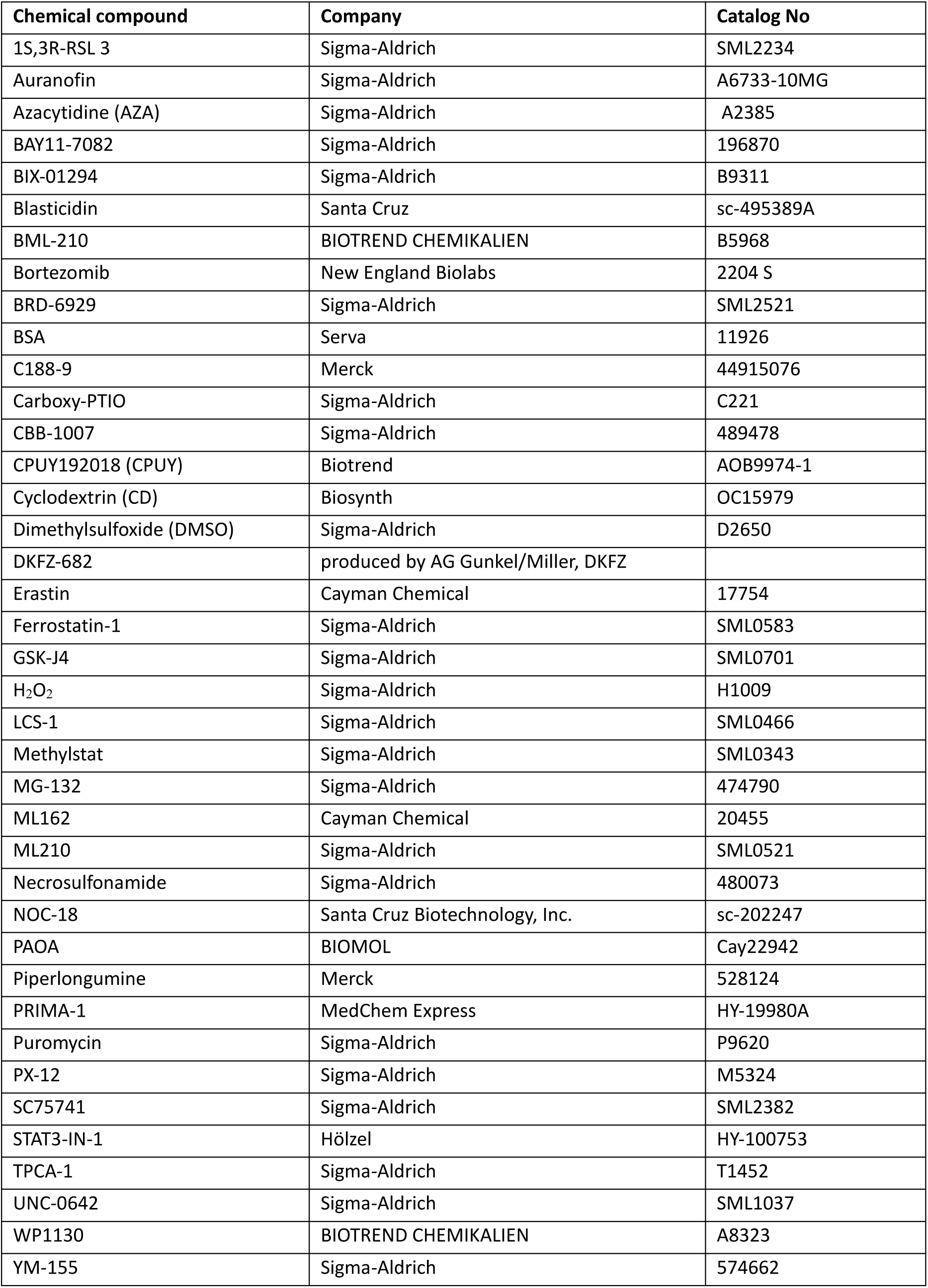
Chemical compounds

**Supplementary Table S2:** Primary and secondary antibodies

| Antigen | Product / Company | Species | Dilution<br>WB |
| --- | --- | --- | --- |
| PRDX1 | #8499/ Cell Signaling | rabbit | 1:1000 |
| PRDX3 | ab128953/ Abcam | rabbit | 1:2000 |
| GCLM | HPA023696/ Sigma-Aldrich | rabbit | 1:100 |
| NRF2 | ab62352/ Abcam | rabbit | 1:1000 |
| xCT/SLC7A11 | 12691/ Cell Signaling | rabbit | 1:1000 |
| GSR | ab124995/ Abcam | rabbit | 1:1000 |
| AKR1C3 | A6229/ Sigma-Aldrich | mouse | 1:500 |
| UGDH | SAB 4503060/ Sigma-Aldrich | rabbit | 1:750 |
| PTGR1 | HPA036724/ | rabbit | 1:250 |
| TXN | ab133524/ Abcam | rabbit | 1:10000 |
| CBR1 | HPA018433/ Atlas Antibodies | rabbit | 1:750 |
| PGD | HPA031314/ Sigma-Aldrich | rabbit | 1:500 |
| BLVRB | HPA041698/ Sigma-Aldrich | rabbit | 1:100 |
| STAT1 | 14994/ Cell Signaling | rabbit | 1:1000 |
| pSTAT (Y701) | 7649/ Cell Signaling | rabbit | 1:1000 |
| STAT3 | 9139/ Cell Signaling | mouse | 1:1000 |
| pSTAT3 (Y705) | 9145/ Cell Signaling | rabbit | 1:1000 |
| STAT5 | 25656/ Cell Signaling | rabbit | 1:1000 |
| pSTAT5 (Y694) | 9359/ Cell Signaling | rabbit | 1:1000 |
| Cas9 | 14697T/ Cell Signaling | mouse | 1:1000 |
| GAPDH | sc-365062/ Santa Cruz | mouse | 1:1000 |
| monoclonal anti β-tubulin | T0198/ Sigma-Aldrich | mouse | 1:1000 |
| IRDye 680LT antimouse IgG | 926-68022/ LI-COR | donkey | 1:5000 |
| RDye 680LT antirabbit IgG | 926-68023/ LI-COR | donkey | 1:5000 |

**Supplementary Table S5:**
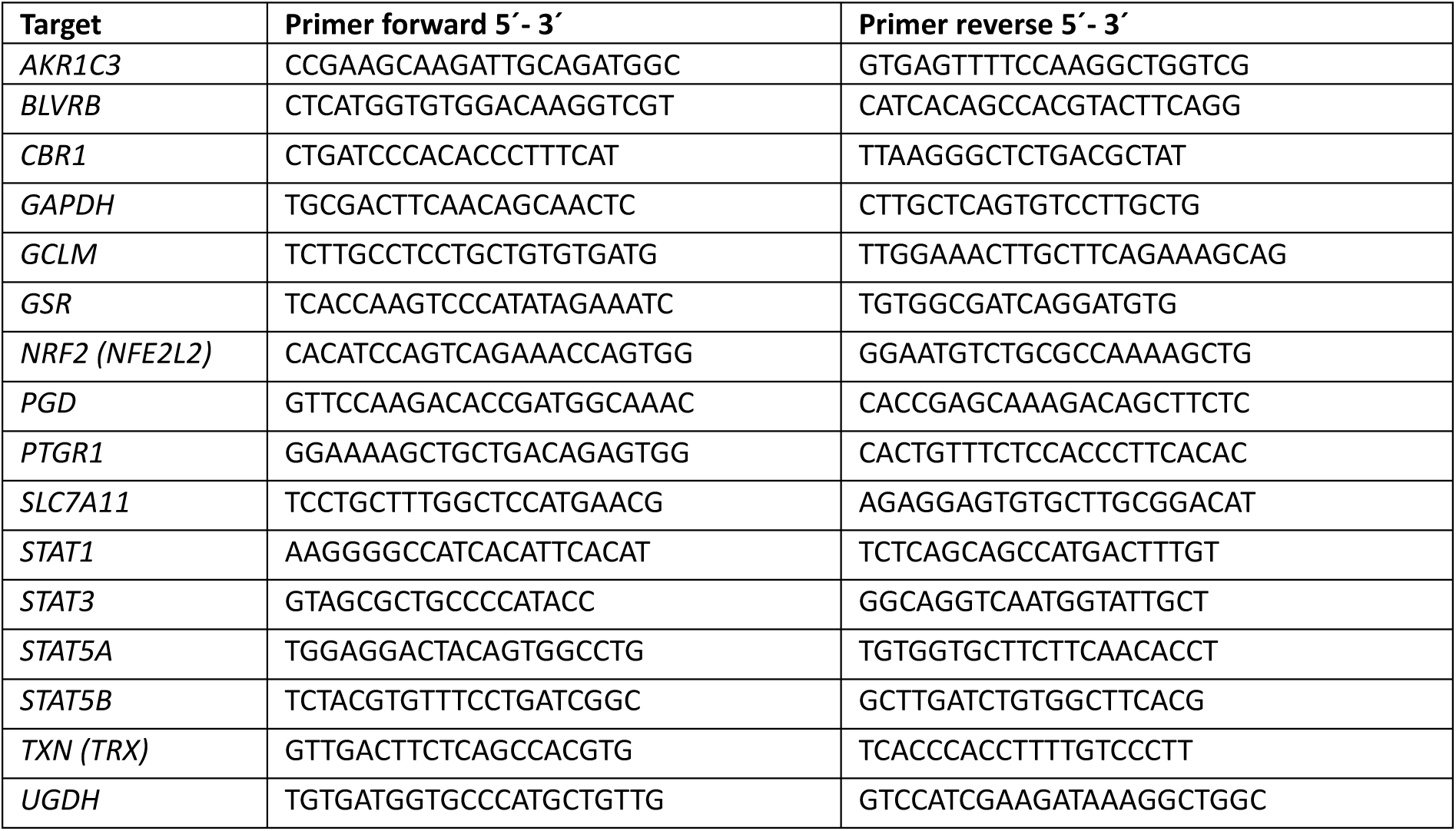
Primer for qPCR

**Supplementary Table S6:**
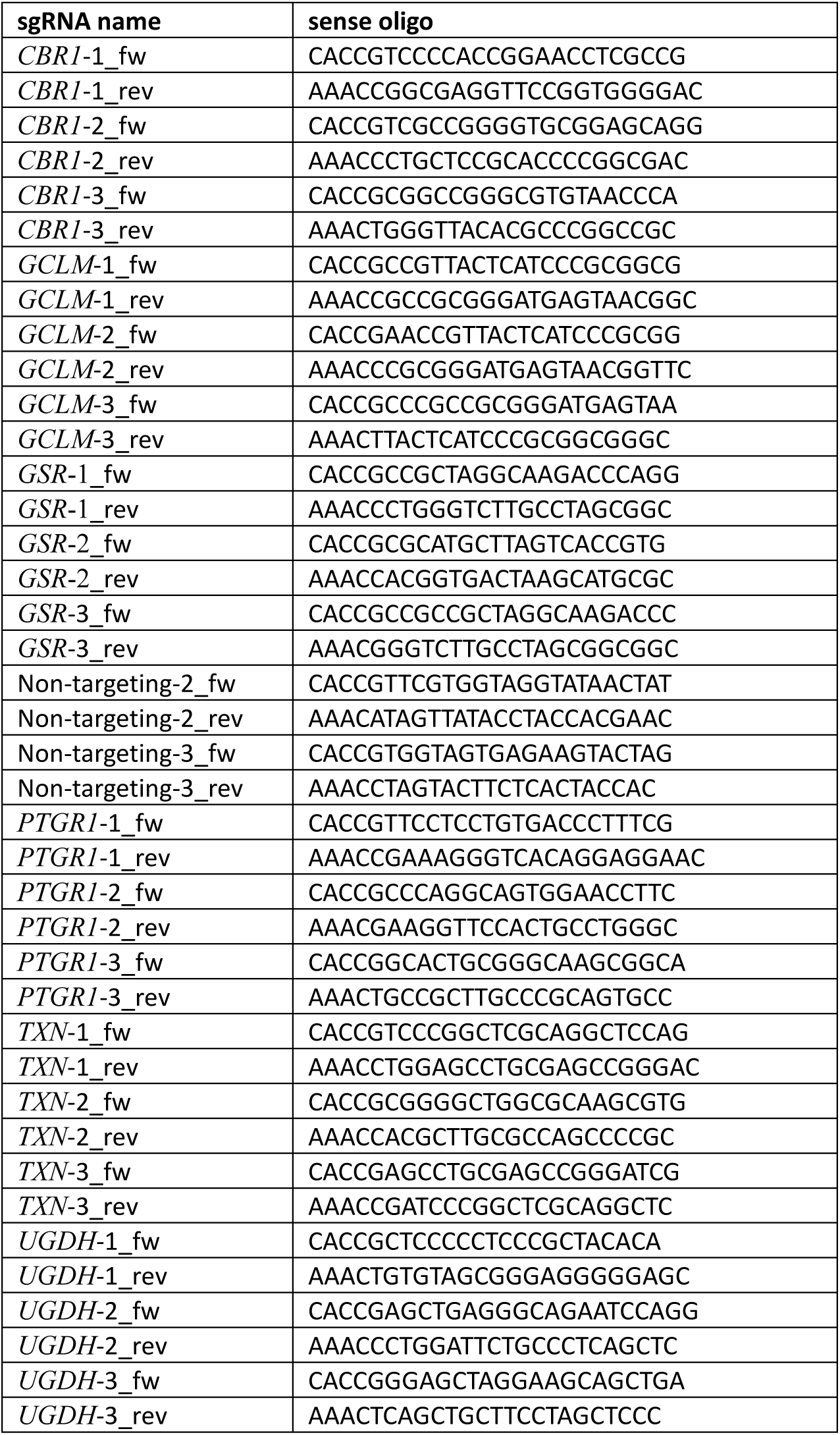
Oligos

